# Alveolar epithelial type 1 cells serve as a cell of origin for lung adenocarcinoma with distinct molecular and phenotypic presentation

**DOI:** 10.1101/2022.10.29.514334

**Authors:** Minxiao Yang, Hua Shen, Per Flodby, Michael D. Koss, Rania Bassiouni, Yixin Liu, Tea Jashashvili, Theresa R. Stueve, Daniel J. Mullen, Amy L. Ryan, John Carpten, Alessandra Castaldi, W. Dean Wallace, Beiyun Zhou, Zea Borok, Crystal N. Marconett

## Abstract

Lung adenocarcinoma (LUAD) is the most common subtype of cancer arising in the distal lung. LUAD encompasses several pathologic subtypes, each with differing clinical outcomes and biological behaviors. However, the molecular and cellular underpinnings of the different subtypes are largely unknown. Understanding which cell populations in the distal lung contribute to LUAD could provide insights into the marked heterogeneity in pathologic features, clinical presentation and responses to therapy of LUAD. Differential expression analysis of lung adenocarcinoma transcriptomes from The Cancer Genome Atlas revealed distinct alveolar epithelial type 1 (AT1) and alveolar epithelial type 2 (AT2) cell signatures within human LUAD with significantly different survival outcomes between tumors expressing AT2 and AT1 gene signatures, suggesting AT1 cells might contribute to a subset of LUAD cases. To address this, we tested the ability of AT1 cells to give rise to LUAD following induction of Kras^G12D^, a known oncogenic driver of human LUAD. Activation of Kras^G12D^ in Gram-domain containing 2 (Gramd2)^+^ AT1 cells gave rise to multiple LUAD lesions, primarily of papillary histology. In contrast, activation of Kras^G12D^ in surfactant protein C (Sftpc^+^) AT2 cells resulted in LUAD lesions of lepidic histology. Immunohistochemistry established that *Gramd2*:Kras^G12D^ lesions were of primary lung origin and not metastatic events. Spatial transcriptomic profiling revealed distinct pathway alterations within Gramd2- and Sftpc-derived LUAD. Immunofluorescence confirmed differences observed in the spatial transcriptomic analysis in expression patterns and distribution of cell-specific markers depending on cell of origin, while universal upregulation of the Krt8 intermediate cell state marker was observed. Our results are consistent with Gramd2^+^ AT1 cells serving as a putative cell of origin for LUAD and suggest that LUAD may be a collection of adenocarcinomas that share a common location within the distal lung but arise from different cells of origin.

## INTRODUCTION

Lung cancer is the leading cause of cancer-related death both within the United States and worldwide (1). Lung adenocarcinoma (LUAD) most often arises in the distal lung and is the most commonly occurring subtype of lung cancer (2). LUAD encompasses several pathologic subtypes, including solid, lepidic, papillary/micropapillary and acinar, each with differing expectations of patient survival, and presents with a wide spectrum of genetic, epigenetic, and pathologic variation (3), the underlying reasons for which are largely unknown. Several studies have linked EGFR status to improved survival outcomes (4), whereas Kras mutations often occur with TP53 co-occurring mutations and result in higher grade LUAD with poorer survival outcomes (5). While it is widely appreciated that different histologic subtypes are associated with differential patient survival outcomes (6), the biology underlying these disparate observations is currently unknown.

LUAD occurrence is localized to the distal lung, where the predominant epithelial cell types are surfactant-producing alveolar epithelial type II (AT2) cells and large, delicate alveolar epithelial type I (AT1) cells (7). AT1 cells cover over 95 percent of the alveolar epithelial surface and are largely responsible for facilitating gas exchange (8). AT1 cells have been thought to be terminally differentiated (9) and consequently unable to proliferate (10). Due to the delicate nature of AT1 cells, they are susceptible to injury, and several mouse lineage studies have shown that AT2, and more recently club cells, are able to repair wounds generated from loss of AT1 cells by proliferating and further differentiating to an AT1 cell identity (11, 12).

The regenerative and self-renewal capacity of AT2 cells helped to establish this cell type as the primary cell of origin for LUAD (2, 13, 14). In contrast, AT1 cells have not been thoroughly explored as a potential cell of origin for LUAD in large part due to a lack of highly cell-specific markers. Outside their distinct morphology, AT1 cells are distinguished by expression of a combination of markers including aquaporin 5 (AQP5), podoplanin (PDPN), homeodomain-only protein homeobox (Hopx), G protein coupled receptor class C group 5 (Gprc5a), and advanced glycosylation end-product specific receptor (human: AGER, mouse: Rage), among others (15).

While each of these has demonstrated utility in identifying AT1 cells, most have modest amounts of RNA expression in other lung cell types, which confounds interpretation of previous attempts to activate oncogenic drivers in AT1 cells. Specific examples of this include Hopx, which has been used previously to demonstrate AT1 to AT2 cell reverse differentiation, as well as the formation of lung nodules (16). However, Hopx mRNA has been observed in AT2 cells in multiple studies (15, 17, 18) raising the possibility that the observed results are due to RNA expression of study-specific drivers in AT2 cells. Studies using the EGFR oncogenic driver in combination with the co-occurring mutations to KEAP1 driven by Gprc5a have shown the ability to form histologically defined LUAD (19); however expression of Gprc5a in subsets of distal basal as well as bronchiolar stem cells (BASC) populations in mice call into question which specific cell type(s) was able to give rise to these LUAD lesions (17). We therefore set out to determine if the AT1 cell can also serve as a cell of origin for LUAD by utilizing bioinformatic analysis in large-scale human LUAD datasets in combination with a mouse model in which oncogenic Kras^G12D^ is activated following tamoxifen inducible Cre recombination selectively in AT1 cells.

## RESULTS

### AT1 cell gene expression signatures are present in human LUAD and predictive of improved patient survival

To comprehensively assess potential AT1 cell involvement in LUAD pathogenesis, we first performed extensive bioinformatic analysis to determine if a gene expression signature for AT1 cells was present within LUAD patient tumors. We hypothesized that LUAD tumors may retain cell-type specific characteristics of the cell of origin from which they arose. This has been observed in multiple tumor types, including breast and prostate (20, 21). Unsupervised hierarchical clustering on the top 10% of differentially expressed genes between LUAD and adjacent non-tumor lung in (AdjNTL) in TCGA LUAD samples, revealed 4 distinct groups of genes which segregated based on overall expression level. These included those with relatively low expression across TCGA LUAD samples (**Figure 1A**, Gene Set 1), moderate to low expression (**Figure 1A**, Gene Set 2), differential expression across the group, with low expression in approximately half of TCGA LUAD samples and high expression in the rest of the cohort (**Figure 1A**, Gene Set 3), and lastly those with high expression overall in LUAD (**Figure 1A**, Gene Set 4). We noted that two distinct groups of gene expression patterns were observed within the TCGA LUAD tumor samples, which we labeled Tumor Cluster A (orange), and Tumor Cluster B (purple).

**Figure 1.**
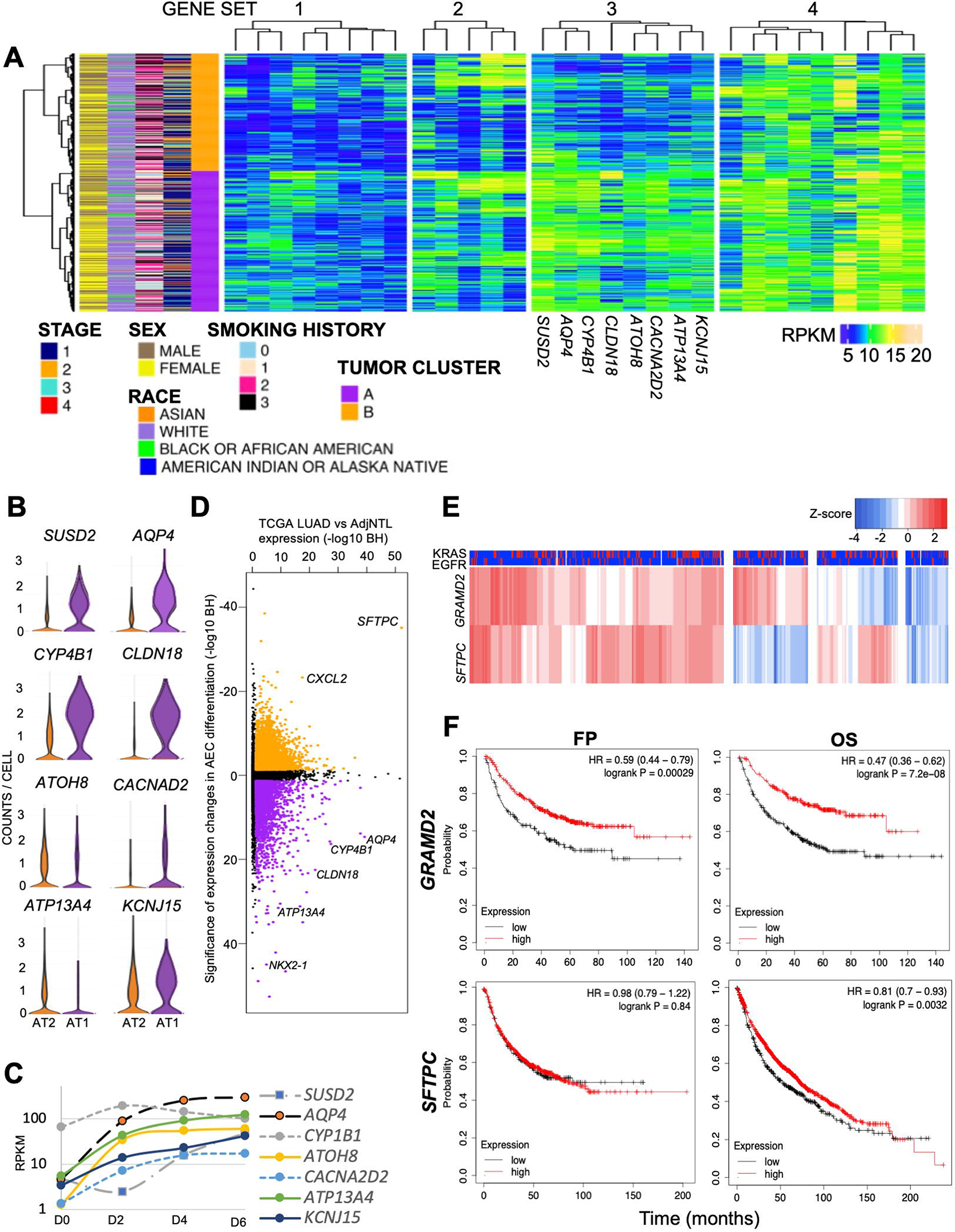
AT1 cell signatures are present within human LUAD and associated with differential survival. A) Unsupervised K-means clustering of top 10% of variant genes within TCGA LUAD tumor samples differentially expressed between LUAD and AdjNTL. Expression levels are denoted by RPKMs (blue = low expression, green = medium expression, yellow/beige = high expression). Patient characteristics are: Sex: yellow = female, brown = male; Race: Asian = orange, White = lilac, Black or African American = green, Indigenous American (denoted American Indian or Alaska Native in TCGA) = blue; Smoking History: 0 = never smoker, 1 = former smoker (>15 years), 2 = former smoker (<15 years), 3 = current smoker; Stage at Diagnosis: IASLC staging criterion. Sample clusters defined by gene expression patterns across the TCGA-LUAD dataset, only tumor samples were included: Tumor Cluster A (purple), Tumor Cluster B (orange). Gene Set 1 (genes with low expression in TCGA-LUAD tumors), Gene Set 2 and 3 (Genes with higher expression in Cluster A, lower expression in Tumor Cluster B), Gene set 4 (Genes with higher expression in both Tumor Cluster A and Tumor Cluster B). B) Violin plot of expression by normal AEC cell type for Gene Set 3 genes (enriched in Tumor Cluster A) from IPF Cell Atlas (ipfcellatlas.com). Y axis = read counts per cell. Orange = normal AT2 cells, purple = normal AT1 cells. C) Expression in RPKMs of Gene Set 3 genes (enriched in Tumor Cluster A) during 2-dimensional (2D) alveolar epithelial cell (AEC) differentiation. X-axis = days in culture; D0 = day 0 (purified AT2 cells), D2 = Day 2, D4 = Day 4 (AEC intermediate cells), D6 = Day 6 (AT1-like cells). D) Significance of changes in transcriptome-wide expression patterns during 2D AEC differentiation (Y-axis) versus significance of changes in transcriptome wide expression patterns between TCGA LUAD and AdjNTL lung. Orange = Gene expression significantly downregulated during 2D AEC differentiation, Purple = gene expression significantly gained during 2D AEC differentiation. Significance is expressed as −log 10 of Benjamini-Hochberg corrected p values. E) Unsupervised hierarchical clustering of TCGA LUAD samples based on *SFTPC* (known AT2 cell markers) and *GRAMD2* (known AT1 cell marker). Row expression is scaled using Z-score (blue = low expression, red = high expression). *EGFR* and/or *KRAS* clinical co-variants are expressed as blue = WT, red = mutated. F) Survival of LUAD patients stratified by expression of *GRAMD2* (known AT1 cell marker) or *SFTPC* (known AT2 cell marker) using KMplot. FP = Free of Progression Survival, OS = overall survival.

To characterize the differential gene expression observed in clusters A and B in more detail, we examined the expression of genes within Gene Set 3 that are normally present within the alveolar epithelium based on previos single cell profiling (17) (ipfcellatlas.com), and showed the most prominent differences in expression between tumor clusters A and B,. We observed that genes from Gene Set 3 were highly enriched for AT1 cell identity (**Figure 1B, Supplemental Figure 1**). To confirm the cell identity associated with the genes expressed in Gene Set 3, we also interrogated their expression in publicly available data on alveolar epithelial cell (AEC) differentiation in 2-dimensional (2D) culture (22), which has previously been shown to recapitulate adult alveolar differentiation that occurs *in vivo* to replace damaged AT1 cells. We observed that all of the genes from Gene Set 3, with the exception of *CYP4B1*, were upregulated during 2D AEC differentiation (**Figure 1C**).

We then asked if AEC-specific signatures were more broadly represented within the TCGA LUAD dataset and observed that there was enrichment for both AT2 (orange) and AT1 (purple) cell signatures in gene expression patterns from the TCGA LUAD data (**Figure 1D**) which is consistent with an alveolar epithelial cell of origin and the location of LUAD tumors in proximity to the distal alveolar space.

Having observed gene expression signatures for both AT2 and AT1 cells in the TCGA LUAD data, we then asked if our observations were a result of the differentiation state of LUAD tumors, where the presence of cell-specific markers was a reflection of a more differentiated tumor state. To determine if the presence of AT1 and AT2 cell signatures was a reflection of differentiation state of the tumor and not cell of origin, we performed clustering on 515 LUAD patient samples present in The Cancer Genome Atlas pan-cancer cohort (23) using *SFTPC*, a known AT2 cell marker, and *GRAMD2*, a known AT1 cell marker (**Figure 1E**). We observed that four distinct clusters emerged, with the dual positive *GRAMD2*^+^ /*SFTPC*^+^ cluster being the largest subgroup; however, there were distinct subgroups where *SFTPC* was expressed and *GRAMD2* was absent, and likewise another group of samples where *GRAMD2* was expressed and *SFTPC* was absent, indicating that distinct cell marker signatures were present in LUAD. We also observed no significant association between tumor stage and cluster identity (**Figure 1A**). Taken together, this suggested that the differentiation state was not driving the presence of cell type specific marker expression in TCGA LUAD tumors.

However, a major feature of differentiated tumors is their improved overall survival. We therefore stratified patients based on the expression of *SFTPC* and *GRAMD2* and evaluated whether their level of expression affected time to recurrence/first progression (FP) using KMplot (24, 25) to determine if, regardless of cell type, the presence of cell-type specific markers was affecting survival (**Figure 1F**). Expression levels of *SFTPC* in lung cancer had no effect on FP (P=0.84), whereas expression of *GRAMD2*^+^ tumors showed significantly longer intervals until FP (p=2.3E-4). Next, we examined patient overall survival (OS) based on expression of alveolar epithelial cell (AEC) cell markers (**Figure 1G**). *GRAMD2* has a large significant protective effect on overall patient survival (P<6.7E-8) whereas the protective effect of *SFTPC* on OS was moderate (p=0.003). Overall, this suggested that the presence of *GRAMD2*, indicative of AT1 cell expression patterns, had a much larger and distinct protective effect on outcomes than *SFTPC*, a known AT2 cell marker. This further strengthened the idea that AT1 cells may be contributing to a subset of human LUAD cases and that the differentiation state of the tumor samples did not explain the variations observed.

### Activation of the oncogenic driver Kras^G12D^ in AT1 cells generates multifocal lung lesions

To functionally test their contribution to LUAD, we activated the Kras^G12D^ oncogenic driver mutation specifically in AT1 cells. Kras mutations contribute to ~30% of LUAD cases, with the G12D variant having a well-established mouse model that can give rise to LUAD when activated in AT2 cells (13, 26). Our laboratory has previously invested in identification of lineage-restricted gene expression patterns for AT1 cell markers within distal lung and identified Gramd2 as highly specific for AT1 cells (15). We subsequently inserted the tamoxifen-inducible CreERT2 transgenic system for genetic recombination into the 3’ end of the endogenous *Gramd2* gene to take advantage of the preexisting regulatory structure that confers AT1 cell specificity to *Gramd2*. *Gramd2-Cre* mice were found to have enriched activation within Aqp5^+^ AT1 cells (27, 28), making it a valuable model for testing if AT1 cells can contribute to LUAD *in vivo*.

*Gramd2*-CreERT2 mice were crossed with Kras^LSL-G12D^ alongside *Sftpc*-CreERT2:Kras^LSL-G12D^ positive mice, which are known to give rise to LUAD derived from AT2 cells. These transgenic models underwent intraperitoneal (IP) injection with optimized tamoxifen (TAM) concentrations, as was previously described (29) and a representative lung from the resultant Kras-activated lungs, hereto named *Gramd2*:Kras^G12D^ or *Sftpc*:Kras^G12D^ were characterized via silication followed by whole lung microCT scanning as performed previously (30) to determine if lesions could form from Kras^G12D^ activation in AT1 cells.

Post-tamoxifen treatment, *Gramd2*:Kras^G12D^ mice formed multifocal lesions throughout the lung (**Figure 2A**), as did the positive control *Sftpc*:Kras^G12D^ mice. No lesions were observed in either wild type C57BL/6, control *Gramd2*-CreERT2 or *Sftpc*-CreERT2 mice with tamoxifen treatment (data not shown), indicating that the Cre promoter drivers alone were unable to drive lesion formation. Corn oil vehicle controls for both *Gramd2*:Kras^LSL-G12D^ and *Sftpc*:Kras^LSL-G12D^ also showed no appreciable lesions (**Figure 2A-B**), indicating that Cre-mediated recombination did not occur unless specifically activated using tamoxifen.

**Figure 2:**
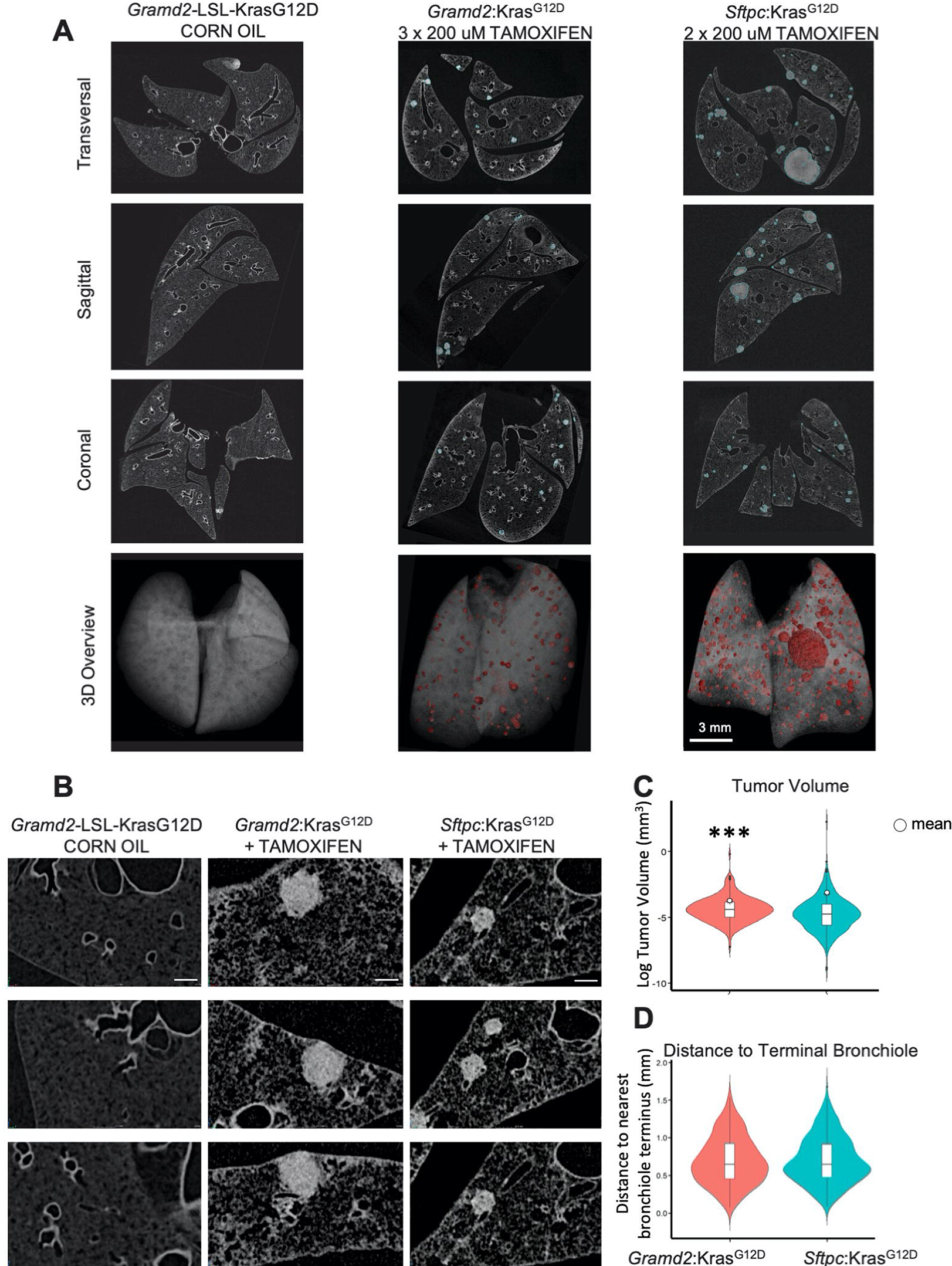
*Gramd2*:Kras^G12D^ mice form hyperplastic lesions throughout the lung at 14 weeks post tamoxifen induction. A) MicroCT scanning was performed on a transgenic mouse lung representative of each genotype, *Gramd2*:Kras^G12D^ (corn oil treated and tamoxifen treated) and *Sftpc*:Kras^G12D^ (tamoxifen treated), at 14 weeks post tamoxifen induction, and the three different sections, transverse, sagittal and coronal, and three-dimensional (3D) overview are listed in order (scale bars in A, 3 mm in each image). B) High magnification scanning was performed on the indicated mouse lungs to identify tumor lesions located in alveoli (scale bars in B, 0.2 mm in each image) ; Significance: (***) = P < 0.001. C) Violin plot indicating the volume of each individual tumor. Teal = *Sftpc*:Kras^G12D^, pink = *Gramd2*:Kras^G12D^. D) Violin plot indicating the distance of each individual tumor to its nearest bronchiole. Teal = *Sftpc*:Kras^G12D^, pink = *Gramd2*:Kras^G12D^. All nodules identified from each representative lung is included.

To quantify the incidence and distribution of lesion formation, we performed a double-blind analysis where lesions were counted, their volumes calculated, and distance from the nearest bronchiole branchpoint measured. We observed approximately half the number of lesions in *Gramd2*:Kras^G12D^ lungs as compared to *Sftpc*:Kras^G12D^ lungs (P < 0.001) which may be due to a number of considerations including but not limited to optimization of Cre activity or pathogenetic properties between cell types. The volume of each individual lesion varied between cell of origin as well, with *Gramd2*:Kras^G12D^ lesions being significantly smaller than those derived from *Sftpc*:Kras^G12D^ (**Figure 2C**). However, the mean distance between the lesion and bronchiole did not differ between *Gramd2*:Kras^G12D^ and *Sftpc*:Kras^G12D^ (**Figure 2D**), which is consistent with the known spatial distribution of AT1 and AT2 cells in the distal alveolar epithelium.

### Gramd2^+^ AT1 cells give rise to LUAD with predominantly papillary histology

To determine the histologic properties of the lesions we observed in *Gramd2*:Kras^G12D^ lungs, matching FFPE sections were stained with hematoxylin and eosin (H&E) and underwent double-blind histologic evaluation alongside *Sftpc*:Kras^G12D^ positive controls. Additional controls evaluated included wild type (WT) C57BL/6, *Sftpc*-CreERT2 Cre driver without Kras^G12D^, *Gramd2*-CreERT2 without Kras^G12D^, and *Kras*-LSL-G12D samples that lacked Cre-drivers. There were no hyperplastic foci detected in H&E sections in negative control sections (**Figure 3A**). AT2 cell driven *Sftpc*:Kras^G12D^ lesions were classified solely as lepidic adenocarcinoma, whereas *Gramd2*:Kras^G12D^ resulted in a mixture of lepidic and papillary adenocarcinoma, with papillary features being the predominant histologic subtype (**Figure 3B**). We also observed papillary adenocarcinoma extending into the bronchial mucosa in *Gramd2*∷Kas^G12D^ lungs, heretofore termed Bronchial Infiltrative Adenocarcinoma (BIA, **Figure 3B**, **Supplemental Figure 2**).These BIAs did not form dense nodules and therefore may have not been detectable using densitometric tracing in the microCT images described above. The identified BIA lesions had classical papillary adenocarcinoma histology within the bronchial lumen but failed to penetrate the lung epithelial lining. This has been observed in multiple mouse models of LUAD (31) but is not a hallmark of human LUAD.

**Figure 3.**
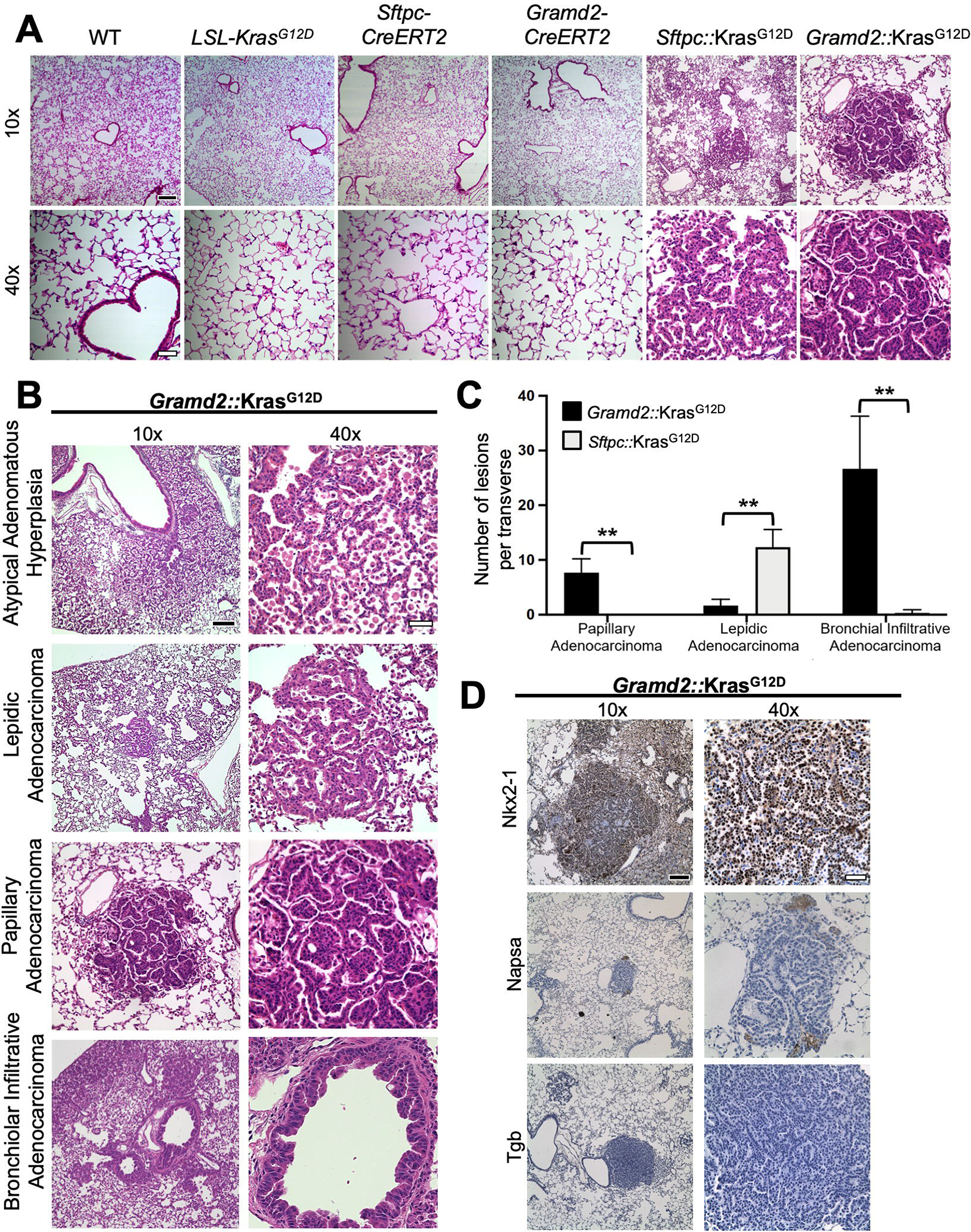
Gramd2^+^ AT1 cells give rise to LUAD with predominantly papillary histology. A) Representative hematoxylin and eosin (H&E) staining of lungs 14 weeks after tamoxifen injection from control mice (C57BL/6 (WT), LSL-Kras-G12D, *Sftpc-*CreERT2, *Gramd2-*CreERT2 as well as *Sftpc*:Kras^G12D^ and *Gramd2*:Kras^G12D^ mice. N = 3 for all genotypes. B) Variation in histologic lesions from *Gramd2*:Kras^G12D^ mouse lungs, including atypical adenomatous hyperplasia (AAH, black arrow), lepidic adenocarcinoma, papillary adenocarcinoma, and bronchial infiltrative adenocarcinoma. (Scale bars in A and B, Black with white outline: 100 μM (10X); white with black outline 25 μM (40X)). N = 3 for all genotypes. C) Quantification of histologic subtypes of lung adenocarcinomas from different mouse genotypes: *Gramd2*:Kras^G12D^ (black bars; n = 3; 14 weeks) vs. *Sftpc*:Kras^G12D^ (grey bars; n = 3; 14 weeks). The bar graphs represent the average number of different types of lesions per section per lung. Data represent means ± SEM. D) Immunohistochemistry (IHC) analysis of *Gramd2*:Kras^G12D^ tumor sections for Nkx2-1, Napsin A, and Thyroglobulin (Tgb). Scale bars: Black with white outline: 100 μM (10X); white with black outline 25 μM (40X).

To quantify differences in histology observed between Kras^G12D^-induced tumors based on cell of origin, a section from each lung was evaluated and the number of lesions of each histologic type recorded from 3 mouse per genotype. This was done to avoid duplicate sampling error that may occur from multiple sections of the same lung. We observed that LUAD of AT2 cell origin was significantly enriched for lepidic histology (P = 0.005). In contrast, LUAD of AT1 cell origin was significantly enriched for papillary histology (P = 0.006) as compared to AT2 cell-initiated tumors (**Figure 3C**). This was intriguing as there are known survival differences between these different histologic subtypes of human LUAD, with lepidic having significantly better overall survival outcomes (32).

To confirm that the adenocarcinomas we observed in *Gramd2*:Kras^G12D^ mice were of lung origin, we performed immunohistochemical staining for known markers of human LUAD which are currently standard of practice for classifying human tumors. Nuclear homeobox protein NKX2-1 (also known as TTF-1 (human), Nkx2-1 (mice)) expression is a known characteristic of lung adenocarcinoma (33, 34). We observed nuclear Nkx2-1 staining within the *Gramd2*:Kras^G12D^ (**Figure 3D**), as well as control *Sftpc*:Kras^G12D^ (**Supplemental Figure 3**), tumors indicating that the tumors were likely primary lung lesions and not metastatic events from other tissues. However, Nkx2-1 can also be indicative of thyroid carcinoma. We therefore stained for Thyroglobulin (Tgb), which is present in thyroid carcinoma but absent in LUAD, to exclude a thyroid origin for these tumors. Tumors from *Gramd2*:Kras^G12D^ and *Sftpc*:Kras^G12D^ were negative for Tgb staining (**Figure 3D, Supplemental Figure 3**). Combined with Nkx2-1^+^ staining, these findings confirm that these adenocarcinomas are likely primary tumors of lung origin. We also performed Napsin A (Napsa) staining to determine if Gramd2+ tumors were enriched for the NAPSA papillary adenocarcinoma marker (35). We observed that *Gramd2*:Kras^G12D^ papillary LUAD derived from AT1 cells were positive for Napsin A staining (**Figure 3D**), whereas AT2 cell derived *Sftpc*:Kras^G12D^ lepidic adenocarcinomas were not (**Supplemental Figure 3**). Additional staining for cytokeratin deposition (Clinical marker AE1/AE3) (46), Cd68 immune infiltration, and Gata3 were also performed to confirm similarities between human LUAD staining patterns and what was observed in *Gramd2*:Kras^G12D^ mice tumors (**Supplemental Figure 3**).

#### Spatial transcriptomic profiling of *Gramd2*:Kras^G12D^ lungs reveals distinct cell-type specific patterning

In order to understand the mechanism(s) by which activation of Kras^G12D^ leads to differential histologic manifestation we undertook spatial transcriptomic profiling using the 10X Genomics Visium platform. Spatial transcriptomics was chosen over other technologies, such as single cell RNA sequencing, to allow integration of histology and spatial distribution with transcriptomic signatures. This was especially important for *Gramd2*:Kras^G12D^ lungs, as multiple histologies were observed including BIA, PA, and LA and the use of spatial technology allowed us to maintain their histologic distinctions throughout downstream analysis. Two biological replicates of *Gramd2*:Kras^G12D^ and *Sftpc*:Kras^G12D^ derived lung sections underwent transcriptomic profiling that included pathologist verified LUAD lesions (**Supplemental Figure 4**). H&E-stained images generated by the Visium pipeline confirmed histologic characterization of these pathologist defined LUAD lesions (**Figure 4A**) and that the *Gramd2*:Kras^G12D^ samples included LUAD of multiple histologies (**Figure 4B**). Expression data from all individual lung samples were then clustered together to evaluate transcriptomic similarities and differences between *Gramd2*:Kras^G12D^ and *Sftpc*:Kras^G12D^ derived lungs. We observed 16 distinct integrated clusters (ICs), so called because each 55 μm spot on the spatial array contains between 1-10 cells (**Figure 4C**).

**Figure 4:**
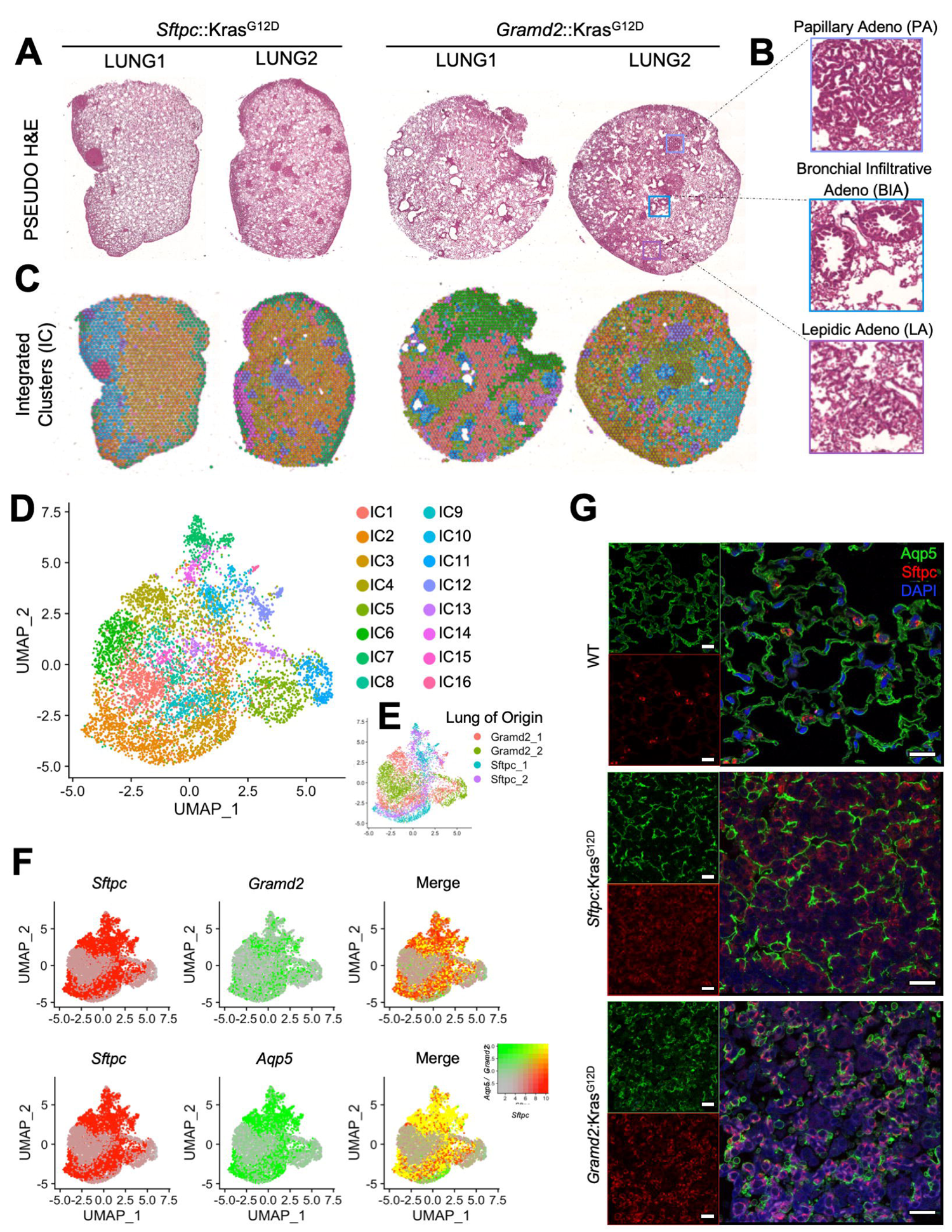
Spatial transcriptomic profiling of *Gramd2*:Kras^G12D^ lungs reveals distinct cell-type specific patterning. A) Pseudo H&E sections generated as part of the Visium 10X spatial transcriptomic profiling performed on 4 lung sections; 2 replicates each *Sftpc*:Kras^G12D^ and *Gramd2*:Kras^G12D^. B) Magnification of distinct histologic regions within *Gramd2*:Kras^G12D^ lung sections. C) Spatial distribution of Integrated Clusters (IC) across all samples within the dataset. Colors indicate distinct ICs. D) UMAP projection of spatial transcriptomic expression data. Colors indicate distinct ICs. E) UMAP projection of spatial transcriptomics expression data colored by sample and replicate of origin. F) UMAP projection indicating degree of overlap between *Sftpc* (AT2) and either *Gramd2* or *Aqp5* (AT1) cell markers. Red = *Sftpc*, green = *Aqp5* or *Gramd2*, yellow = co-expression of *Sftpc* and either *Aqp5* or *Gramd2* within the 10X array spot indicated. G) Immunofluorescence imaging of colocalization between Sftpc (red) and Aqp5 (green) within either wild type C57BL/6 (WT), N = 3 per genotype. Scale (white bar) = 10 μM (63x).

Uniform manifold approximation and projection (UMAP) was used for dimensional reduction and visualization of IC similarities and differences (**Figure 4D**). We observed a high degree of concordance between lung sections irrespective of sample identity, with the notable exceptions of IC5 and IC11 which were enriched in *Gramd2*:Kras^G12D^ samples and IC7, IC14, and IC16 which were enriched in *Sftpc*:Kras^G12D^ samples (**Figure 4E**).

To further interrogate the differences between *Gramd2*:Kras^G12D^ and *Sftpc*:Kras^G12D^ derived LUAD, we examined the distribution of *Sftpc* and *Gramd2* within all samples. It was anticipated that *Gramd2*:Kras^G12D^ and *Sftpc*:Kras^G12D^ derived LUAD would have mutually exclusive cell-type marker expression, with Gramd2-driven LUAD enriched for AT1 cell markers, and Sftpc-driven LUAD enriched for AT2 cell markers. Instead, we observed that both Sftpc- and Gramd2-derived LUAD were enriched for both cell type markers (**Figure 4F**).

We then wanted to determine if the dual expression of both AT1 and AT2 cell markers that we observed in the majority of human LUAD transcriptomic profiling (**Figure 1D**), was a result of cell mixtures within spatial transcriptomic spots, or if instead the cells lose distinct marker expression as part of the carcinogenic process. To do this, we performed immunofluorescent staining for Gramd2 and Sftpc on both *Gramd2*:Kras^G12D^ and *Sftpc*:Kras^G12D^ derived LUAD (**Figure 4G**). Immunofluorescence staining confirmed that both Sftpc and Gramd2 were expressed in LUAD derived from both *Sftpc*:Kras^G12D^ and *Gramd2*:Kras^G12D^, but interestingly the staining was observed to be mutually exclusive, indicating that the LUAD lesions were composed of multiple cell types. Intracellular staining patterns for Sftpc were similar in both *Sftpc*:Kras^G12D^ and *Gramd2*:Kras^G12D^ derived LUAD, with Sftpc staining surrounding the periphery of cuboidal cells reminiscent of AT2 cell architecture. In contrast, Aqp5 staining patterns varied dramatically depending on cell of origin. In *Sftpc*:Kras^G12D^, Aqp5 staining was long and planar reminiscent of normal AT1 architecture and occurred at the boundaries of the Sftpc+ hyperproliferative regions. In contrast, Aqp5 staining in *Gramd2*:Kras^G12D^ derived LUAD was markedly rounded and dispersed throughout the tumor. Taken together, these results suggested that the contribution to differential histology observed between *Sftpc*:Kras^G12D^ and *Gramd2*:Kras^G12D^ derived LUAD may be in part aided by the disruption and redistribution of AT1 cell architecture.

#### Expression analysis reveals distinct transcriptomic signatures between *Gramd2*:Kras^G12D^ and *Sftpc*:Kras^G12D^ LUAD

We then sought to determine if the histologically defined LUAD lesions recapitulated known gene expression signatures associated with human LUAD. To do this, we applied ESTIMATE, a tool developed on pan-cancer signatures from TCGA to estimate tumor content from expression data (36) (**Figure 5A**). We observed that ESTIMATE computed and histologically defined LUAD array spots were significantly enriched in histologically defined LUAD clusters, but not completely concordant as many spots included in histologically defined LUAD clusters did not meet the tumor purity threshold of ≥ 0.85 (**Figure 5B**). UMAP projection demonstrated that spots with high tumor purity segregated into three distinct regions (**Figure 5C**), corresponding to BIA, LA, and PA histopathology. Closer examination of these high tumor purity regions demonstrated that multiple clusters were included in both the LA (IC12 and IC16), and PA (IC4) histologies (**Figure 5D**). The identity of IC14 was less clear in histologic definition, as the lepidic adenocarcinoma presented with some micropapillary features and was interspersed between more classically defined PA and LA histologies. The exception was IC11, which corresponded precisely to the BIA phenotype. This demonstrated that both PA and LA histologies could be defined by their transcriptomic signatures. It also revealed that there was a distinct difference between the BIA and PA histology. These clusters also corresponded to cell-of-origin (**Figure 5E**), as the PA and BIA histologies were exclusively observed in *Gramd2*:Kras^G12D^ and the LA histology was observed in both *Gramd2*:Kras^G12D^ and *Sftpc*:Kras^G12D^ derived LUAD.

**Figure 5:**
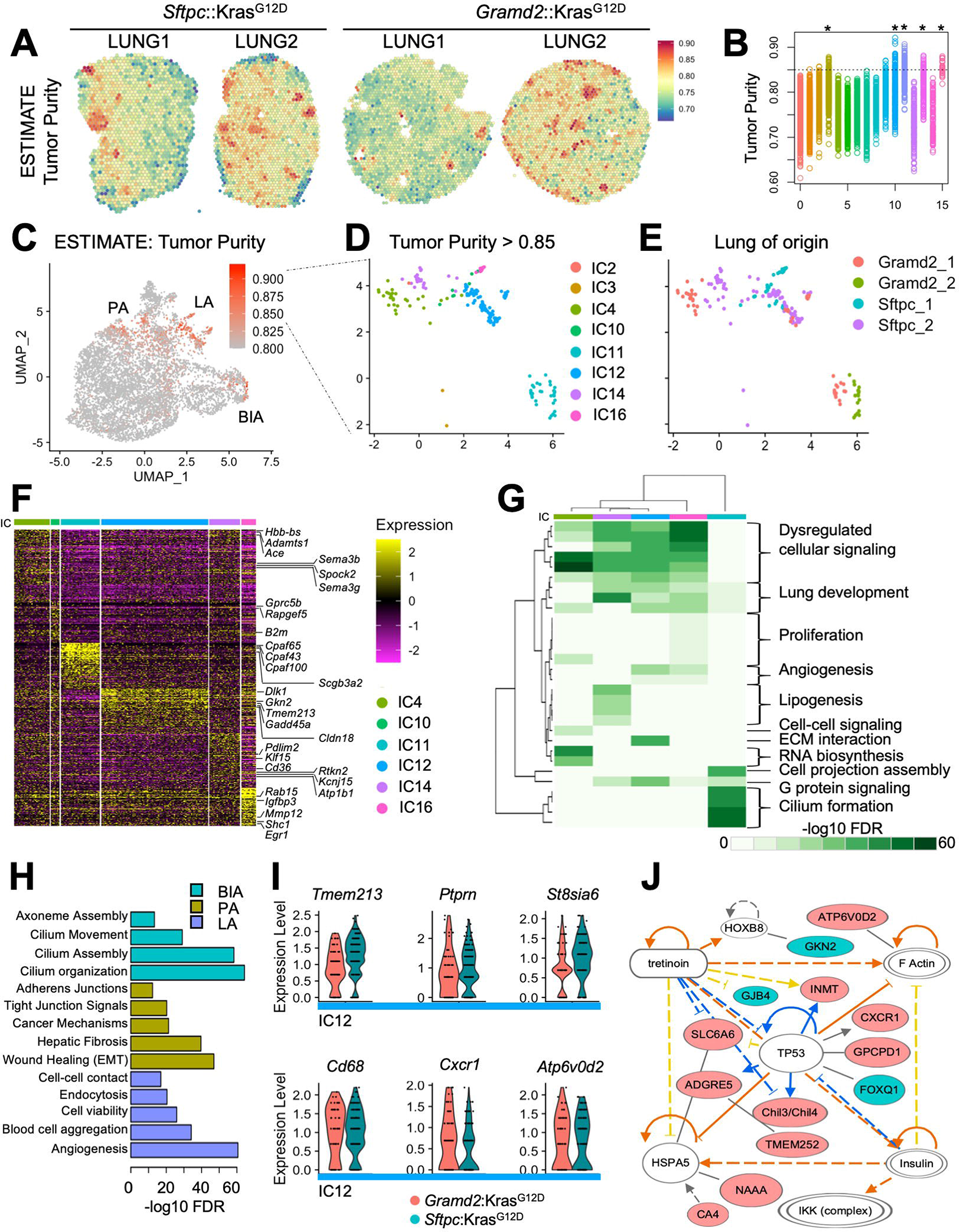
ESTIMATE analysis of high tumor purity reveals distinct transcriptomic signatures between *Gramd2*:Kras^G12D^ and *Sftpc*:Kras^G12D^ LUAD. A) ESTIMATE was applied to quantitate tumor purity within samples; Red = high tumor purity, blue = low tumor purity. B) Correlation between tumor purity and cluster identity. Colors = Integrated Cluster ID C) UMAP projection of expression data from all samples colored by tumor purity; Red = high tumor purity, gray = low tumor purity. D) UMAP projection of a subset of spatial transcriptomics spots with tumor purity greater than or equal to 0.85. Colors = Integrated Cluster ID, Teal (IC11) = bronchiolar infiltrative adenocarcinoma (BIA) histology, blue + pink (IC 12+16) = lepidic adenocarcinoma, purple = lepidic adenocarcinoma with or without co-occurring micropapillary features, green (IC4) = papillary adenocarcinoma (PA) histology. E) UMAP projection of spots with tumor purity ≥ 0.85. Colors indicate sample origin. F) Heatmap of top 200 differentially expressed genes between ICs from spots with ≥ 0.85 tumor purity. Only ICs with >5 occurrences were included to minimize technical biases. Select genes relating to carcinogenesis or lung cell type-specific markers are indicated. Yellow = highly expressed, purple = little to no expression. Integrated Cluster (IC) color as indicated previously. G) Heatmap of Panther gene set enrichment analysis (GSEA). The −log10 of false-discovery rate (FDR) corrected P values are indicated; dark green = highly significant FDR of Gene Ontology (GO) category enrichment, white = non-significant GO enrichment. H) IPA analysis of differential activity in separate histological patterns. Purple = lepidic adenocarcinoma (LA), peridot = papillary adenocarcinoma (PA), and teal = bronchiolar infiltrative adenocarcinoma (BIA). I) Violin plots of top differentially expressed genes between *Sftpc*:Kras^G12D^ (teal) and *Gramd2*:Kras^G12D^ (pink). J) Ingenuity pathways analysis of differentially expressed genes between *Sftpc*:Kras^G12D^ (teal) and *Gramd2*:Kras^G12D^ (pink) within IC12. White = no difference in gene expression between cell of origin, orange lines = positive effect, blue lines = inhibitory effect, yellow line = gene expression relationships displaying opposite regulatory relationships to predictive models.

In order to determine transcriptional differences that were driving the segregation of clusters based on histology, we undertook differential expression analysis among those ICs with at least 5 occurrences among all samples measured. Supervised clustering between clusters of the top 200 differentially expressed genes revealed distinct expression patterns unique to each cluster (**Figure 5F**). Included among these were several known lung cell type specific markers; including the previously described AT1 cell markers *Sema3b*, *Spock2*, *Sema3g*, and *Rtkn2* (15, 37, 38). Interestingly, genes that were enriched in tumor cluster A in human TCGA LUAD (**Figure 1A**), such as *Kcnj15*, were also significantly enriched in ICs of high tumor purity supportive of a role for AT1 cells in human LUAD. Of note, IC11 was enriched for several *Cpaf* genes, including but not limited to *Cpaf65*, *Cpaf43* and *Cpaf100*, all of which have distinct functions in the formation and assembly of ciliated structures (39), indicating that the BIA phenotype not observed in human LUAD is likely derived from low-level Gramd2 expression in ciliated cells.

To further determine how known carcinogenic genes varied within each IC, we performed gene set enrichment analysis (GSEA) using the Protein ANalysis THrough Evolutionary Relationships (PANTHER) (40, 41). GSEA showed that several pathways with distinct functions in carcinogenesis were significantly enriched, including proliferation, dysregulated cellular signaling, disruptions in cell to cell signaling and extracellular matrix deposition (**Figure 5G**). Of note, angiogenesis was observed to be highly enriched in IC12 and IC16 of LA histology, whereas multiple dysregulated cellular signaling pathways were observed in IC4 associated with classical PA. To understand exactly which signaling pathways were dysregulated, we performed Ingenuity Pathways Analysis (IPA) on PA (IC4), BIA (IC11), and LA (IC12 and IC16) enriched genes (**Figure 5H**) which takes into account functional relationship and known direct and indirect signaling networks to predict common activation patterns and upstream regulators. We observed significant enrichment of VEGF-mediated angiogenesis pathway enrichment in LA, whereas PA was enriched for TGF beta-mediated hepatic fibrosis and EMT pathways. Of note, IPA pathways analysis confirmed that IC11 enriched in BIA histology was enriched for cilia formation, motility and axoneme assembly, suggesting that there is a subset of ciliated cells present in the lower bronchiolar space with limited *Gramd2* expression that was not detected by previous methods.

One of the striking features of *Gramd2*:Kras^G12D^ LUAD lesions was that multiple histologic subtypes were observed during H&E analysis (**Figure 3**), including LA as well as PA. As LA was observed in both *Gramd2*:Kras^G12D^ and *Sftpc*:Kras^G12D^ LUAD lesions, we sought to determine if there were significant gene expression differences within LA between the two cells of origin. Comparison of Sftpc- vs Gramd2- driven enrichment within IC12 LA observed in both cells of origin revealed 90 differentially expressed genes. Genes elevated in *Sftpc2*:Kras^G12D^ LUAD included *Tmem213*, *Ptprn*, and *St8sia6* (**Figure 5I**), whereas multiple genes involved in immune cell function including *Cd68* and *Cxcr1* were elevated in LA derived from *Gramd2*:Kras^G12D^. Indeed, a comprehensive analysis of ESTIMATE-defined immune cell enrichment across all tissue sections showed marked changes in immune cell composition between *Gramd2*:Kras^G12D^ and *Sftpc*:Kras^G12D^ lung sections (**Supplemental Figure S5**). IPA analysis of these 90 genes revealed a common axis centered around the central *TP53* pathway (**Figure 5J**). This was striking, as *TP53* loss occurs significantly with Kras oncogenic activation in LUAD and is significantly associated with response to immunotherapy (42), further suggesting our model is an accurate recapitulation of human LUAD carcinogenesis. Stromal cell composition also varied between *Gramd2*:Kras^G12D^ and *Sftpc*:Kras^G12D^ LUAD, with enrichment of alpha-SMA (*Acta2*) observed in IC13 solely derived from *Gramd2*:Kras^G12D^ LUAD (**Supplemental Figure S6)**. LUAD derived from both *Sftpc*:Kras^G12D^ and *Gramd2*:Kras^G12D^ also exhibited elevated proliferation rates that were similar between *Gramd2*:Kras^G12D^ and *Sftpc*:Kras^G12D^ LUAD, suggesting that the ability to proliferate when an oncogenic driver mutation was present did not vary between the two cell types **(Supplemental Figure S7)**.

#### *Gramd2*:Kras^G12D^ and *Sftpc*:Kras^G12D^ derived LUAD both exhibit Krt8^+^ intermediate cell states

One of the top differentially expressed genes between histologically defined LUAD clusters and adjacent normal lung was elevated expression of *Krt8* (**Figure 6A**). While LUAD of BIA histology (IC11) had one of the highest levels of Krt8 expression (**Figure 6B**), elevated Krt8 was observed across all ICs with high tumor purity (**Figure 6C**). Indeed, the concordance between *Krt8* expression and tumor purity was extremely significant (R = 0 .41, P < 2.1E-16). This is intriguing, as Krt8 is a known marker of stress-induced transitional cell states in mouse lung (12, 43). We therefore sought to investigate the relationship between Krt8 and histologic presentation in Gramd2- and Sftpc-driven LUAD.

**Figure 6:**
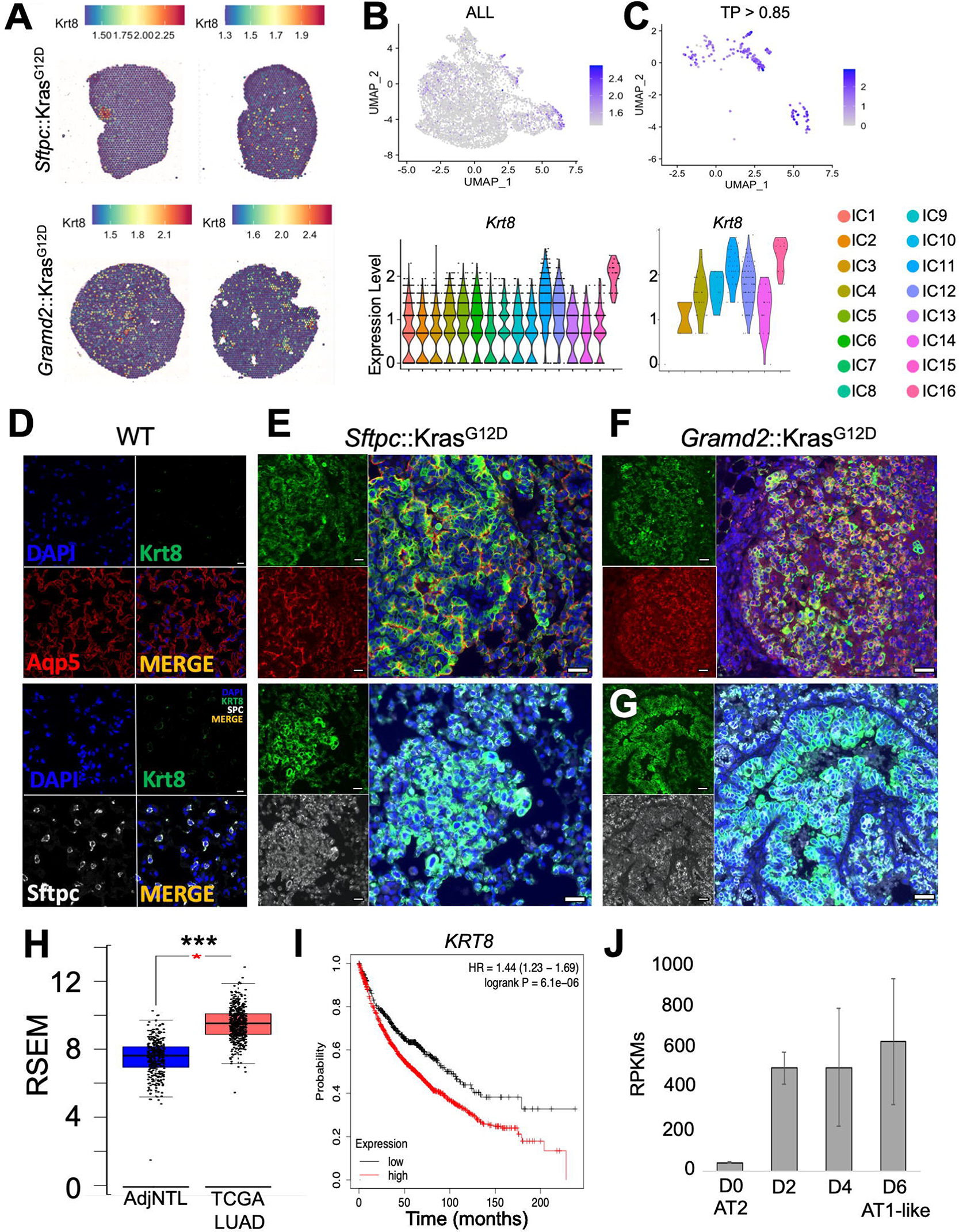
Krt8 is elevated in LUAD and upregulated during normal AEC differentiation and in both *Gramd2*:Kras^G12D^- and *Sftpc*:Kras^G12D^-derived LUAD. A) Spatial distribution of *Krt8* distribution throughout the entire dataset. Red = high *Krt8* expression, blue = low *Krt8* expression. B) UMAP projection of IC integration in all array spots within the dataset (top) and violin plot of *Krt8* expression within individual ICs in all array spots (bottom). Colors indicate distinct ICs. *Krt8* expression, grey = low *Krt8* expression and violin plot of *Krt8* expression within individual ICs in array spots with tumor purity of ≥ 0.85 (bottom). Colors indicate distinct ICs. D) IF staining of Krt8^+^ in control B57J/L mice lungs. Red = Gramd2, white = Sftpc, green = Krt8, blue = DAPI, orange = merged. Sftpc was originally captured in TRITC and was converted to greyscale in ImageJ. E) IF of *Sftpc*:Kras^G12D^ mouse tumors, colors are as in (D). F) *Gramd2*:Kras^G12D^ LUAD with lepidic patterning, colors are as in (D). G) *Gramd2*:Kras^G12D^ LUAD with bronchiolar-infiltrative (BIA-LUAD) phenotype, colors are as in (D). White scale bar in lower right corners = 20 μM (20X). H) Expression of Krt8 in TCGA LUAD. Blue = adjacent non-tumor lung (AdjNTL), red = LUAD. I) Overall Survival (OS) of LUAD patients stratified by Krt8 expression. Plot generated in Kmplot (33, 34). Black = low expression, red = high expression. J) Expression of Krt8 in publicly available AEC 2D culture bulk RNAseq (22). Expression measured in reads per kilobase of gene per millions mapped in sample (RPKMs).

To confirm our transcriptomic analysis results that *Krt8* was elevated in both Gramd2- and Sftpc-driven LUAD we evaluated Krt8 staining patterns in the Kras^G12D^ initiated mouse tumors by performing immunofluorescence on both *Gramd2*:Kras^G12D^ and *Sftpc*:Kras^G12D^ derived tumors and compared the distribution of Krt8 staining patterns. Staining patterns in control C57BL/6 mice were consistent with previous literature (12) (**Supplemental Figure S8**), in that the alveolar epithelium expressed little to no Krt8^+^ under normal physiologic conditions (**Figure 6D**). Also consistent with previous literature was the presence of Krt8^+^ cells within the bronchial tree (**Supplemental Figure S9**). We then examined how the distribution of Krt8 varied between *Sftpc*:Kras^G12D^-derived lepidic LUAD and *Gramd2*:Kras^G12D^-derived papillary LUAD and observed that Krt8 staining was elevated in both tumor types (**Figure 6E-F**). Krt8 deposition in both tumor types surrounded tumor nuclei.

We then sought to determine if the tumor nuclei surrounded by Krt8+ signal showed cell-type specific marker expression consistent with the cell in which Kras^G12D^ had been activated. Co-staining of Krt8 and Sftpc in *Sftpc*:Kras^G12D^ lepidic LUAD showed that tumor nuclei were surrounded by Sftpc expression, as anticipated (**Figure 6E**). Unfortunately, the Gramd2 antibody was incompatible with Krt8, therefore Aqp5 was used as a surrogate AT1 cell marker to interrogate AT1 cell distribution in *Sftpc*:Kras^G12D^ lepidic LUAD. Aqp5 was also present in *Sftpc*:Kras^G12D^ as observed previously (Figure 4G), and costained with Krt8 (**Figure 6F**). *Gramd2*:Kras^G12D^ lepidic LUAD also showed distinct overlap between Aqp5 and Krt8 staining, consistent with increased Krt8 expression in LUAD arising from both cells of origin. Bronchiolar-infiltrative LUAD observed in *Gramd2*:Kras^G12D^ also showed distinct expression of Krt8 surrounding tumor nuclei which were also positive for Sftpc (**Figure 6G, Supplemental Figures 2-3**). Our results indicated that Krt8 upregulation was present in LUAD tumors from both AEC cells of origin, and may be reflective of the carcinogenic stress response in distal lung.

Krt8 expression levels are associated with poor outcomes in human LUAD (44). Expression of KRT8 was elevated in human LUAD from the TCGA-LUAD dataset (**Figure 6H**), and we also observed elevated KRT8 expression was associated with poorer OS in human LUAD (**Figure 6I**) in Kmplot (33, 34). This was intriguing, as Krt8 has been described as a transitional cell state during the maladaptive repair phase following fibrosis and may play a role in the carcinogenic process. We then evaluated whether KRT8 levels are induced during stress conditions in human AEC differentiation of purified human AT2 cells in 2D culture and evaluating KRT8 expression over the course of differentiation into AT1-like cells as previously described (45). We observed that KRT8 was elevated during human AEC differentiation (**Figure 6J**).

In sum, we observed Krt8 expression in lesions derived from both models, but interestingly and unexpectedly, we also observed that LUAD in both models stained for both AT1 and AT2 cell markers, suggesting that activation of the Kras^G12D^ oncogenic driver in both *Gramd2*:Kras^G12D^ and *Sftpc*:Kras^G12D^ result in Krt8^+^ intermediate cell states. We therefore hypothesize that either AT1 or AT2 cells, when exposed to stress conditions, such as activation of Kras^G12D^, adopt a defensive transitional cell state to attempt to resolve the injury. This Krt8^+^ cell state, which has been observed following bleomycin injury to resolve into reformation of the normal alveolar epithelium is unable to do so when subject to genetic alterations and continues to progress toward the formation of hyperplastic foci that retain the cellular memory of the cell type from which they originated.

## DISCUSSION

Results of this study suggest that Gramd2^+^ AT1 cells can give rise to histologically verified LUAD, and therefore that Gramd2^+^ AT1 cells are able to serve as a cell origin of LUAD. Furthermore, different cells of origin can give rise to LUAD with different histologic patterns. Gramd2+ AT1 cells gave rise primarily to papillary adenocarcinoma, whereas Sftpc^+^ AT2 cells gave rise solely to lepidic adenocarcinoma. Lepidic adenocarcinoma is known to have improved overall survival outcome compared to all other types of LUAD (~92% 5-year survival rate), and papillary adenocarcinoma also has improved outcomes (~70% 5-year) compared to acinar and solid LUAD histologies (<15% 5-year). Our results suggest that other mature epithelial cell types resident in the distal lung may also contribute to LUAD that manifests in different histologic patterns not observed in our model systems. It has been previously suggested that the bronchiolar stem cell (BASC) resident at the bronchoalveolar junction (BADJ) can also give rise to LUAD (46), though a thorough characterization of histologic subtypes has to our knowledge not yet been performed. It is also possible that the distal respiratory epithelium, which is present in humans but not in mice, may harbor unique cell populations that can give rise to LUAD.

Gramd2 was initially identified as an AT1 cell marker in 2D culture of AT2 cells differentiating into AT1-like cells whose activation was conserved across human, mouse, and rat (15). This initial study saw induction of Gramd2 after 4 days in culture, when other known AT1 markers were also activated, such as Aqp5, Pdpn, and Ager (among others). However, by Day 4 in culture there was an obvious induction of the intermediate cell marker KRT8 in this cell population, which could indicate that Gramd2, as has been observed for Hopx (47), is activated in both the AEC-intermediate cell state, which is present at basal levels in the adult lung (47), and in the fully differentiated “mature” AT1 cell. Igfbp2 is currently considered the most lineage restricted marker for mature AT1 cells (48), therefore it would be interesting to see if Kras^G12D^ activation within the Igfbp2-restricted AT1 cell population was able to give rise to papillary LUAD as well.

Our results also indicate that both AT1 and AT2 cell lineages activate Krt8 expression. Krt8^+^ cells have been identified as stress-induced intermediate cells that can arise from both AT2 and the more proximally located Club cells that subsequently differentiate into AT1 cells and repopulate the lung epithelial lining (12). To our knowledge, this study is the first demonstration that LUAD may progress through this intermediate state as well, but unlike bleomycin-induced fibrosis the cells are unable to resolve the initial stressor and continue to grow out of this intermediate population. Elevated expression of KRT8 is known to predict poor patient outcome for LUAD and is associated with epithelial-to-mesenchymal transition (44), further implicating this intermediate transitional cell state in the etiology of LUAD.

Another unexpected finding in Gramd2^+^ AT1-derived LUAD in mice was the development of Bronchiolar Invasive Adenocarcinoma (BIA). This has been observed previously in Gprc5a^+^ cell-specific induction of EGFR oncogenic activity (31) and has largely been characterized as a species-specific phenomenon, as this does not occur within human LUAD. These species-specific effects may be due to the lack of distal respiratory epithelium and the interactions between cells of origin and these regions. Additionally, in human LUAD, Kras^G12D^ mutations are significantly associated with concurrent loss of TP53 function (TP53-LOH). In transgenic mouse models combination of Kras+/− with TP53-LOH increases the multiplicity, size and degree of dedifferentiation observed in tumors (26). It is possible that a combination of Kras^G12D^ with TP53 LOH in Gramd2^+^ AT1 cells may result in higher grade tumors, reflecting known histologic patterns observed in human LUAD with mixed histology.

Based on our spatial transcriptomic analysis, the BIA phenotype was derived from a ciliated cell population, which were histologically, spatially, and transcriptomically distinct from the alveolar papillary lesions we observed. BIA was also highly enriched for cilia formation, axoneme assembly, and other pathways with known involvement in multiciliated cell function. Our microCT scanning results demonstrated that nodule formation was restricted to bronchiole regions and equidistant from terminal bronchi with Sftpc-driven lesions. This would argue that either BIA microCT was not able to detect the BIA phenotype as it did not grow in distinct nodules, or that these were equidistant from the terminal bronchi and derived from a population of ciliated cells in near proximity to the alveolar space. Taken together, it appears that the BIA lesions, derived from ciliated cells and not consistent with any known histology in human LUAD, do not serve as a cell of origin in human LUAD.

Pathway analysis comparison between lepidic and papillary adenocarcinoma revealed distinct differences, despite all instances being driven by the same Kras^G12D^ mutation. Lepidic adenocarcinoma was enriched for Vegf-mediated angiogenic pathways, whereas TGF-beta mediated EMT was enriched in papillary adenocarcinomas. It will be interesting to see if promising small molecular inhibitors that have been highly successful in preclinical studies targeting these pathways that subsequently failed in clinical trials may have had muted effects due to being tested on LUAD derived from different cells of origin. We also observed distinct differences in immune and stromal cell composition depending on of origin. *Gramd2*:Kras^G12D^ showed upregulation of alpha-SMA (*Acta2*), which has been associated with EMT. In contrast, *Sftpc*:Kras^G12D^ was associated with increased PD-L1 (Cd274), among other factors. It will be interesting to see if cell of origin affects response to immunotherapy in future studies.

Taken together, our results demonstrate that AT1 cells can serve as a cell of origin for LUAD. Additionally, our results indicate that different cells of origin in the distal give rise to LUAD of different histology and molecular phenotype. This work is consistent with recent observations that Hopx^+^ AT1 cells have proliferative capacity (49). Importantly, we found that the same mutation, Kras^G12D^, caused different cellular signaling disruptions depending on the cell in which the oncogene was activated, which could lead to the diverse phenotypic presentations observed. Our work suggests that LUAD is not one cancer type that arises solely from AT2 cells but instead is a collection of cancers that occur within spatial proximity in the distal lung, each with distinct morphologic, phenotypic, and transcriptomic differences that derive from the cell of origin in which the cancer arises.

## MATERIALS AND METHODS

### TCGA Bioinformatic Analysis

For initial differential gene expression calculations used in Figure 1A, 515 TCGA LUAD Level 3 RPKM expression data were filtered based on differential expression between LUAD and AdjNTL samples to include the top 10% variant genes across the dataset. Samples were clustered using K-means and cut using CutTree and visualized using heatmap.2 in R (version 3.2.11). For additional interrogation specifically for GRAMD2 and *SFTPC*, GENCODE v26-annotated, log2 FPKM-UQ gene expression values from 453 TCGA LUAD samples were visualized using the heatmap.3 function (https://github.com/obigriffith/biostar-tutorials/blob/master/Heatmaps/heatmap.3.R). EGFR and KRAS mutational data were acquired using cBioportal (50, 51). Expression values were transformed into Z-scores and samples were clustered in an unsupervised manner.

### Kaplan Meier Survival Analysis

Survival analysis was performed on a compilation of multiple cohorts of lung cancer present in Kmplot whose expression was profiled using Affymetrix geneChip Arrays which were composited and normalized within the Kmplot functionality (Kmplot.com) (24, 25). Data were split based on lower quartile expression (top ¾ of patients vs. lower ¼ of patients) and significance calculated independently for progression free-survival (FP), and overall survival (OS). Significance with Kmplot is calculated using the logrank P value.

### Generation of *Gramd2*-CreERT2 transgenic mouse and mouse models

The *Gramd2*-creERT2 knockin mouse line was produced by Applied Stem Cell, Inc. (Milpitas, CA). A CRISPR/Cas-assisted gene targeting approach was used in mouse ES cells to knockin creERT2 into the endogenous *Gramd2* locus. Homologous recombination in ES cells (derived from a 129 mouse sub-strain) was assisted by simultaneous electroporation of the targeting vector together with Cas9 and a guide RNA (gRNA) that directed Cas9 to a site immediately upstream of the stop codon of *Gramd2*. Detailed characterization of this mouse is described in a separate manuscript. The *Sftpc*-CreERT2 mouse line has been used previously to track AT2 cell fate (65, 66). *Gramd2*-CreERT2 mice were crossed to heterozygous *Kras*^LSL-G12D^ mice (Jackson laboratory, Stock: #008179). *Gramd2*-CreERT2 and *Sftpc*-CreERT2 mice were in a B6/129 mixed strain background (129 N1), while Kras^LSL-G12D^ mice are in a B6/129S4 mixed strain background (C57BL/6 N10).

### Mouse Genotyping

Pups were weaned before the 21^st^ day after birth. For each pup, 0.1 cm of tail was cut and stored in a 1.5 mL microcentrifuge tube. 100 μL DirectPCR (#102-T, Viagen) with 10 μL proteinase K (50 μg/ml, Qiagen, DNeasy Blood & Tissue Kit, #69504) was then added to each tube and incubated with the tail overnight at 60°C. To inactivate the proteinase K, a two-hour incubation at 92°C was performed the next day, and crude DNA was ready for use. A concentration of 50 ng/ul was used as a template for subsequent genotyping PCR reactions. PCR was performed using GoTaq G2 Hot Start Polymerase (Promega, Lot: 0000362382). For *Gramd2* gene, reactions consisted of 1 μL DNA, 5 μL buffer, 2 μL MgCl2, 0.5 μL dNTP, 1 μL common forward primer, mutant reverse primer and wild type reverse primer, 0.12 μL polymerase, and nuclease-free water to a total volume of 20 μL. For the *Kras* gene, the PCR recipe is similar except using 3 μL of DNA template. For *Sftpc* gene, the PCR recipe is similar except for two sets of primers: one for the mutant *Sftpc* allele and wild type *Sftpc* allele. For *Gramd2* and *Kras* genes, the reaction was incubated for 2 minutes at 94°C, 35 cycles including 20 seconds at 94°C, 30 seconds at 58°C and 90 seconds at 72°C, and 10 minutes at 72°C using the C1000 Touch Thermal Cycler (Bio-Rad, Los Angeles, CA). For *Sftpc* gene, the reaction was incubated for 3 minutes at 94°C, 35 cycles including 30 seconds at 94°C, 30 seconds at 58°C and 40 seconds at 72°C, and 2 minutes at 72°C using the C1000 Touch Thermal Cycler (Bio-Rad, Los Angeles, CA). Primer sequences are listed in the Supplemental Methods.

### Gel Electrophoresis

PCR products were loaded into a 2.0% agarose gel, which was prepared using LE Agarose (VWR, LIFE SCIENCE, CAS: 9012-36-6) and nuclease-free water. The PowerPac Basic Power Supply (BIO-RAD, PowerPac Basic) was used to perform gel electrophoresis at 80V for 1 hour. Gels were imaged on a Molecular Imager ChemiDox XRS+ (BIO-RAD, Los Angeles, CA).

### Tamoxifen Administration

Tamoxifen (Sigma-Aldrich, CAS: 10540-29-1) powder was dissolved in corn oil (Sigma-Aldrich, CAS: 8001-30-7) to a final concentration of 40 mg/mL. To ensure that tamoxifen was fully dissolved in corn oil, the solid-liquid mixture was gently shaken and incubated in a shaking incubator (SHEL LAB, model: SSI5R; cat: #33-804R) at 65°C for 2 hours. Six-week-old mice were injected with tamoxifen via intraperitoneal injection. For *Gramd2*:Kras^LSL-G12D^, *Gramd2*-CreERT2, Kras^LSL-G12D^ and WT mice strains, a stock concentration of 40 mg/kg tamoxifen with a final dosage of 200 mg/kg per mouse was introduced via intraperitoneal (IP) injection three times on alternating days. For *Sftpc*:Kras^LSL-G12D^ and *Sftpc*-CreERT2 mice strains, the stock concentration of 40 mg/mL tamoxifen was administered by IP injection twice on alternating days to a final dosage of 100 mg/kg.

### Dissection and Processing of Lung Samples

At 14 weeks post-tamoxifen injection, mice were euthanized by injecting 100 μL Euthasol (ANADA #200-071, Virbac) via intraperitoneal injection. The lungs were then perfused with phosphate buffered saline (PBS, 21-031-CV, CORNING) and inflated with 4% paraformaldehyde (PFA, CAS: 30520-89-4, Sigma-Aldrich) at a pressure of 25 cm water. After fixation, the lungs were transferred into a 50 mL Falcon tube and fixed in 4% PFA overnight. Fixed lungs were then washed three times with 15 mL sterile PBS and stored in 70% ethanol for future use.

### Embedding and Sectioning

Embedding and sectioning were accomplished with assistance of the USC Translational Research Core in the USC School of Pharmacy. Briefly, lung samples were processed in an automatic tissue processor (Thermo Scientific Spin Tissue Processor Microm STP 120) following a standard gradient of dehydration (70%, 80%, 95% and 100% of ethanol), clearing (Clear-Rite 3) and paraffin infiltration. After embedding, samples were sectioned at 5 μm using a rotary microtome (Thermo Fisher Microm HM310 Rotary Microtome) and affixed to clean slides.

### Silication & Micro CT Scanning

After fixation with 4% PFA using insufflation, the lungs of *Gramd2*-*CreERT2*; *Kras*LSL-G12D and *Sftpc*-*CreERT2*; *Kras*LSL-G12D mice were gently rinsed with PBS, and then with 50% ethanol, before placing in 70% ethanol. This rinse was repeated three times. Next, lung samples were dehydrated using an ethanol gradient at room temperature, in 70%, 80%, and 90% ethanol solution each for 2 hours, and then in 100% ethanol, left overnight. After lung samples were dehydrated using ethanol, we used the low-surface-tension solvent hexamethyldisilazane (HMDS) (SHBG4111V, Sigma-Aldrich) to incubate samples. For infiltration, the lung samples in this study were left in the chemical hood for 1-2 hours until they were thoroughly dried and solidified as previously described (52).

The dried and solidified lungs underwent micro-CT scanning at 6-8 μm voxel resolution at the USC Molecular Imaging Center in the Department of Radiology at Keck School of Medicine, USC using An GE Phoenix nanotom *M* micro-CT scanner system. The scans were performed using the following parameters: 60kVp, 200 μA using 1440 projections along a 360-degree rotation at one frame per second rate. Raw image data were reconstructed into 16-bit DICOM images. Visualization of x-ray image data, 3D surfaces renderings of lesions inside of lungs, and quantification of lesion volumes and distances between the lesions and terminal bronchioles were performed using VGSTUDIO MAX 3.3.2.170119 64 bit (© Copyright 1997-2019 by Volume Graphics GmbH).

The micro-CT images and 3D surface renderings were used to segment the multifocal lesions throughout the lung, calculate each lesion’s volume (mm3), and collect the x, y, and z coordinates of the center point for each lesion. We also collected the x, y, and z coordinates of landmarks placed in each terminal bronchiole just before the alveolar ducts. Then all-possible distances between each lesion and landmark of each terminal bronchiole were calculated, and minimal distance was chosen as the distance between the lesion and terminal bronchiole. This calculation was repeated until we found all possible minimal distances between the lesion and terminal bronchiole.

### H&E Staining

H&E staining was accomplished with assistance of the USC Immunohistochemistry laboratory, Department of Pathology, Keck School of Medicine, USC. The slides were deparaffinized and stained in Hematoxylin and Eosin using an automated stainer (Varistain™ Gemini ES Automated Slide Stainer).

### IHC Staining

Immunohistochemistry (IHC) staining was accomplished with assistance of the USC Immunohistochemistry laboratory in the Department of Pathology at the Keck School of Medicine, USC. First, sections were baked at 60°C for 1-2 hours and then left to cool for fifteen minutes. After cooling, the following steps were processed in the BOND-III Fully Automated IHC/ISH Staining System (Leica BioSystems). Sections were first deparaffinized and then underwent antigen retrieval by incubating with EDTA-based epitope retrieval solution (BOND Epitope Retrieval Solution 2, Cas#AR9640, Leica BioSystems) for 20 minutes. After antigen retrieval, slides were washed with distilled water 3 times for 2 minutes each. Next, processed sections were incubated with Multi-Cytokeratin (AE1/AE3) (10 mg/mL, CAS#PA0909, Leicabiosystems), Anti-CD68 Monoclonal Antibody, Unconjugated, Clone 514H12 (37 mg/L, CAS#PA0286, Leicabiosystems), Anti-GATA-3 monoclonal Antibody, Clone L50-823 (1:20, CAS#CM405B, BIOCARE), Anti-Napsin A monoclonal Antibody, Clone TMU-Ad02 (1:100, CAS#CM388A, BIOCARE), Anti-thyroglobulin monoclonal Antibody, Clone 2H11 + 6E1 (1:250, CAS#340M-15, Cell Marque), and TTF-1 (1:500, Cell Marque, Cas#343M-96) Antibody for fifteen minutes. After primary incubation, sections were washed with 0.2% Tween diluted in PBS for two minutes for three times at 25°C. Next, sections were incubated with post primary block and polymer (BOND IHC Polymer Detection Kit (DS9800), Leica micro-biosystem). Each incubation lasted for eight minutes. After that, sections were blocked with peroxide for five minutes, then staining was performed with DAB Chromogen (BOND IHC Polymer Detection Kit (DS9800), Leica micro-biosystem) for ten minutes. In addition, sections were stained with Mayer’s Hematoxylin (American Master Tech). The processed sections were then taken out of the Leica Bond III Auto-stainer and were dehydrated by dipping in 95% isopropyl alcohol for one minute, 100% isopropyl alcohol for one minute, and xylene for one minute. Dehydrated sections were mounted and covered with glass coverslips.

### Visium 10X Spatial Transcriptomic Profiling

Spatial transcriptomic profiling was performed at the Molecular Genomics Core, a part of the Norris Comprehensive Cancer Center and using the manufacturer’s instructions. Briefly, two biological replicates (one block for each) of *Gramd2*:Kras^G12D^ and *Sftpc*:Kras^G12D^ underwent initial 10 μM sectioning at the Tissue Pathology Core, part of the Norris Comprehensive Cancer Center. Sections were H&E stained (as above) and underwent pathology review. Regions of Interest (ROIs) then underwent sample preparation including test slide sample sequencing using the Visium Tissue Section Test Slide (PN-2000460, 10X Genomics, Dublin, CA, USA). RNA quality assessment of test slides was determined by first extracting RNA using the Qiagen RNeasy FFPE kit (#73504, Qiagen, Hilden, Germany) and measuring RNA concentration using a Qubit fluorometer (#Q33238, ThermoFisher Scientific, Waltham, MA, USA). The percentage of total fragments >200nt in length was determined by running the samples using a 4200 Tapestation (#G2991AA, Agilent, Santa Clara, CA). Once samples were determined to be of high enough quality to continue, 5 μM FFPE sections were then placed in a 42°C water bath and once rehydrated adhered to the Visium Spatial Gene Expression Slide (PN-2000233, 10X Genomics). Subsequently, samples were dried at 42°C for 3 hrs in a desiccation chamber and underwent deparaffinization using Qiagen Deparaffinization Solution (Cat # 19093) at 60C for 2 hr. Subsequently H&E staining (#MHS16, #HT110116 Millipore Sigma, Burlington MA, USA) was performed according to Visium technical parameters (CG000409) and images were captured with a Zeiss Axioscan2 microscope using a 10x objective. Decrosslinking was performed according to 10X standard protocol (CG000407) and immediately hybridized to the Visium Mouse Transcriptome probe set V1.0, which contained 20,551 genes targeted by 20,873 probes. Post-probe extension, sequencing library construction was performed using unique sample indices using the Dual Index Kit TS, Set A (PN-1000251) for Illumina-compatible paired-end sequencing (2×100). Raw sequencing data was processed with the Space Ranger pipelines (10x Genomics) spaceranger mkfastq for demultiplexing of sequencing data, and spaceranger count for alignment and unique molecular identifier counting, and tissue detection and alignment.

### Data Accessibility

The spatial transcriptomic data generated in this study are publicly available in the Gene Expression Omnibus (GEO) as GSE215858. OS and PFS data are available through KMplot (http://kmplot.com/lung/) (24, 25). TCGA data is available for download through the Gene (GDC) portal (https://portal.gdc.cancer.gov/).

### Bioinformatic Analysis of Spatial Transcriptomic Data

Assay spot count data generated by CellRanger was imported into R (v 4.0.5) and analyzed using R Studio (v 1.4.1717). Seurat (v4.1.1) (53) was used for the majority of analysis. Individual samples underwent SC transform normalization of data (54), clustering, and principal component analysis to define variable features before merger of all lung samples into one dataset. Principal component analysis was then performed on all merged assay spots, and IC cluster identities were assigned. Cell cycle scoring was adapted from Seurat after conversion of preprogrammed human to mouse IDs using the mouse genome database (68). ESTIMATE analysis was performed using the ESTIMATE R package (v1.0.13)(55) by converting human gene IDs present in the ESTIMATE package to their mouse orthologs. Where multiple mouse orthologs existed for one human gene, all orthologs were included. ESTIMATE analysis of tumor purity, immune score, and stromal score was then added as Metadata to the Seurat object containing all lung section samples for integrated analysis. Pearson correlation between two variables within the Seurat object was done using the cor.test() function in base R. IPA (v 01-20-04) was performed on differentially expressed genes between *Sftpc*:Kras^G12D^ and *Gramd2*:Kras^G12D^ in IC12 using both direct and indirect interactors to generate network analysis ranked by the total number of interactors as well as differentially expressed genes based on histologic patterning across sample types.

### EdU Incorporation and Detection

For EdU labeling, mice underwent intraperitoneal injection (IP) with 5-ethynyl-2`-deoxyuridine (EdU) (component A, Invitrogen™ Click-iT™ EdU Cell Proliferation Kit for Imaging, Cat# C10337) first dissolved in DMSO (component C) to a stock concentration of 50 mM, then subsequently diluted in PBS (Gibco™, PBS pH 7.4, Cat#10010023) to a final concentration of 20 mM prior to injection. EdU detection was performed per the manufacturer’s instructions with 500 μL of Click-iT® reaction cocktail ((Invitrogen, Cat# C10337)) per coverslip. For 1 sample reaction, the following amounts of the kit components were mixed in 430 μl 1X Click-iT® reaction buffer (component D): 50 μl buffer additive (component F, previously diluted in deionized water and kept frozen in small aliquots), 20 μl copper (II) sulfate solution (Component E, 100 mM CuSO4) and 1.2 μl Alexa Fluor 488 azide (Component B, previously dissolved in, component C, 70 μl DMSO).

### Immunofluorescence

Primary antibodies used for co-staining were rabbit-anti-Gramd2 (1:100, ATLAS biological, CAS #HPA 029435), rabbit-anti-Sftpc (1:300, Seven Hills, CAS#WRAB-9337), and Rat-anti-Keratin, type II/ Cytokeratin 8 antibody (1:100, DSHB, CAS#ab531826). Antibody signal of Cytokeratin 8 was amplified using biotinylated anti-rat antibody (1:300, 0.5 mg, FisherScientific Vector Laboratories, CAS#NC9016344), followed by staining with Streptavidin, Alexa Fluor™ 488 conjugate (1: 300, Invitrogen, CAS#S11223). Antibody signal of Gramd2 or Sftpc was visualized by staining with IgG (H+L) Highly Cross-Adsorbed Goat anti-Rabbit, Alexa Fluor™ 594 (1:300, FisherScientific Vector Laboratories, CAS#A11037)

### Statistical Analysis

Initial tumor identification studies were performed on a cohort of 5 mice per genotype. Follow-up studies were performed on groups with ≥ 3 mice per genotype. Statistical comparisons between two groups were made using a Student’s two-tailed t-test. Comparisons between multiple groups were performed using ANOVA with Bonferroni multiple testing correction. Two biological replicates of *Gramd2*:Kras^G12D^ and *Sftpc*:Kras^G12D^ underwent spatial transcriptomic profiling. Differential expression analysis in spatial data were made using Seurat with the FindMarkers functionality (54).

## Acknowledgements

The authors would like to thank Junji Watanabe, PhD and the Translational Research Lab in the School of Pharmacy for assistance with tissue embedding and sectioning, Seth Ruffins, PhD and the Optical Imaging Core at the Broad Center for Regenerative Medicine and Stem Cell Research (CIRM) for assistance with immunofluorescence, Peter Conti, M.D, PhD and the Molecular Imaging Center for assistance with microCT, and David Wesley Craig, PhD and the Molecular Genomics Core. Core usage at the Norris Comprehensive Cancer Center is supported by the NCI (P30 CA014089). Funding for this study was provided by the American Cancer Society through a Research Scholar Grant (RSG-20-135-01) to CNM, R35 HL135747 to ZB, the Wright Foundation, and the Departments of Surgery and of Translational Genomics, Keck School of Medicine, USC.

