## Supplemental Methods & Figures for "Alveolar epithelial type 1 cells serve as a cell of origin for lung adenocarcinoma with distinct molecular and phenotypic presentation"

### SUPPLEMENTARY MATERIALS & METHODS

Primer sequences for mouse genotyping:
***Gramd2*-CreERT2** [forms a product from reverse primer specificity with either mutant (Cre-cassette containing) or WT]

*Gramd2*-CreERT2 common forward: 5’- CTAGTCCTGTCCTCGTCCTATC-3’,

*Gramd2*-CreERT2 mutant allele reverse: 5’- GGGAAACCATTTCCGGTTATTC-3’,

*Gramd2*-CreERT2 WT allele reverse: 5’- CACATCCCAGCCTTCTCAAA-3’

***Sftpc*-CreERT2** (uses separate primers for WT and CreERT2 amplicons)

*Sftpc*-CreERT2 WT allele forward: 5’- TGGTTCCGAGTCCGATTCTTC-3’,

*Sftpc*-CreERT2 WT allele reverse: 5’- CCTTTTGCTCTGTTCCCCATTA-3’,

*Sftpc*-CreERT2 mutant allele forward: 5’- TGAGGTTCGCAAGAACCTGATGGA-3’,

*Sftpc*-CreERT2 mutant allele reverse: 5’- ACCAGCTTGCATGATCTCCGGTAT-3’,

***Kras-LSL-G12D*** [Forms a product from forward primer specificity with either mutant (Cre-cassette containing) or WT]

*Kras*$^{LSL\text{-}G12D}$ common reverse: 5’- CTGCATAGTACGCTATACCCTGT-3’,

*Kras*$^{LSL\text{-}G12D}$ mutant forward: 5’- GCAGGTCGAGGGACCTAATA-3’,

*Kras*$^{LSL\text{-}G12D}$ WT forward: 5’- TGTCTTTCCCCAGCACAGT-3’,

### SUPPLEMENTAL FIGURES:

**Figure S1**

scRNAseq counts per gene per cell (Ipfcellatlas.com)

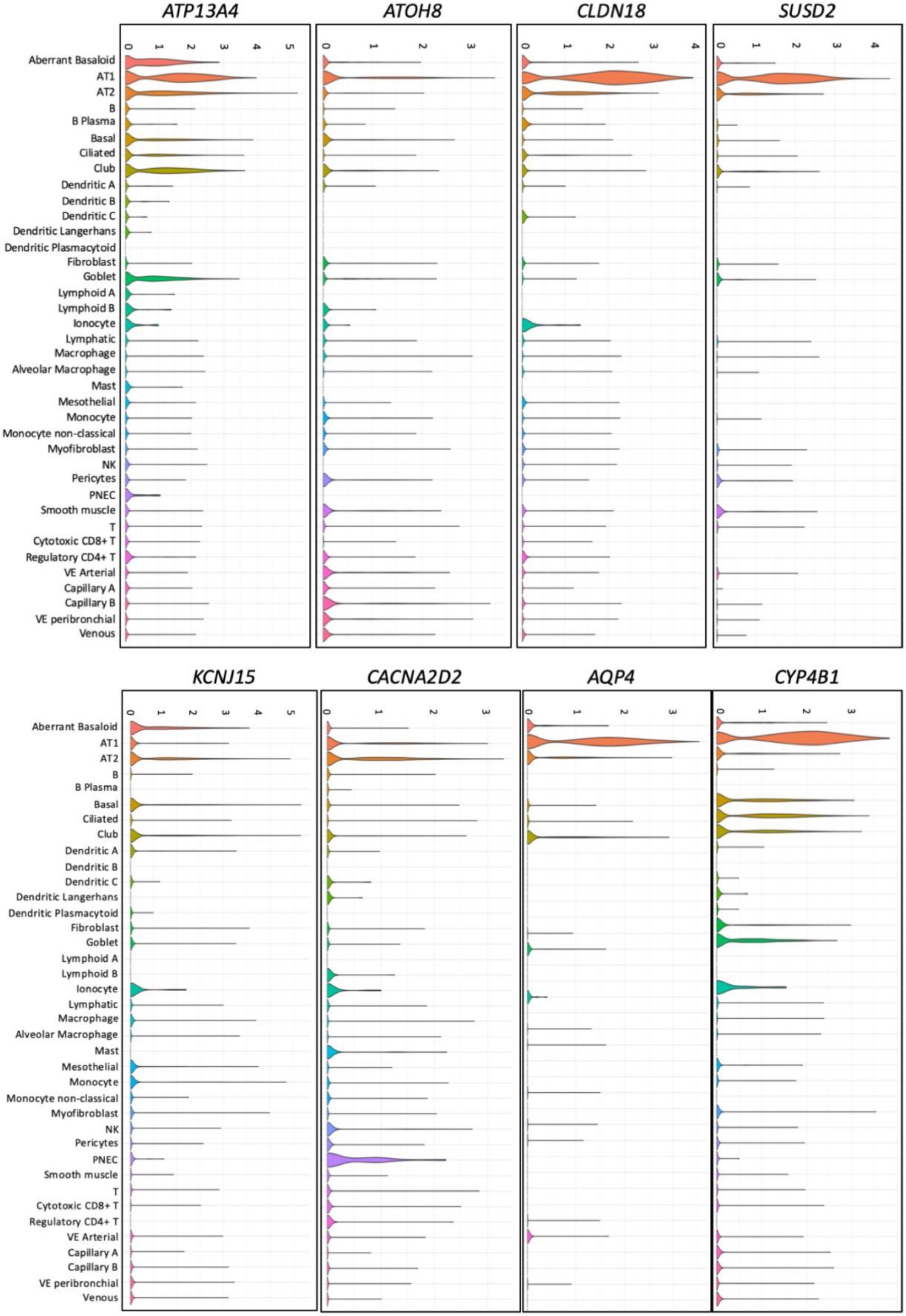

**Supplemental Figure 1: Cell type specific expression of TCGA LUAD Gene Set 3 genes in the IPFCellAtlas**

(17).

**Figure S2**

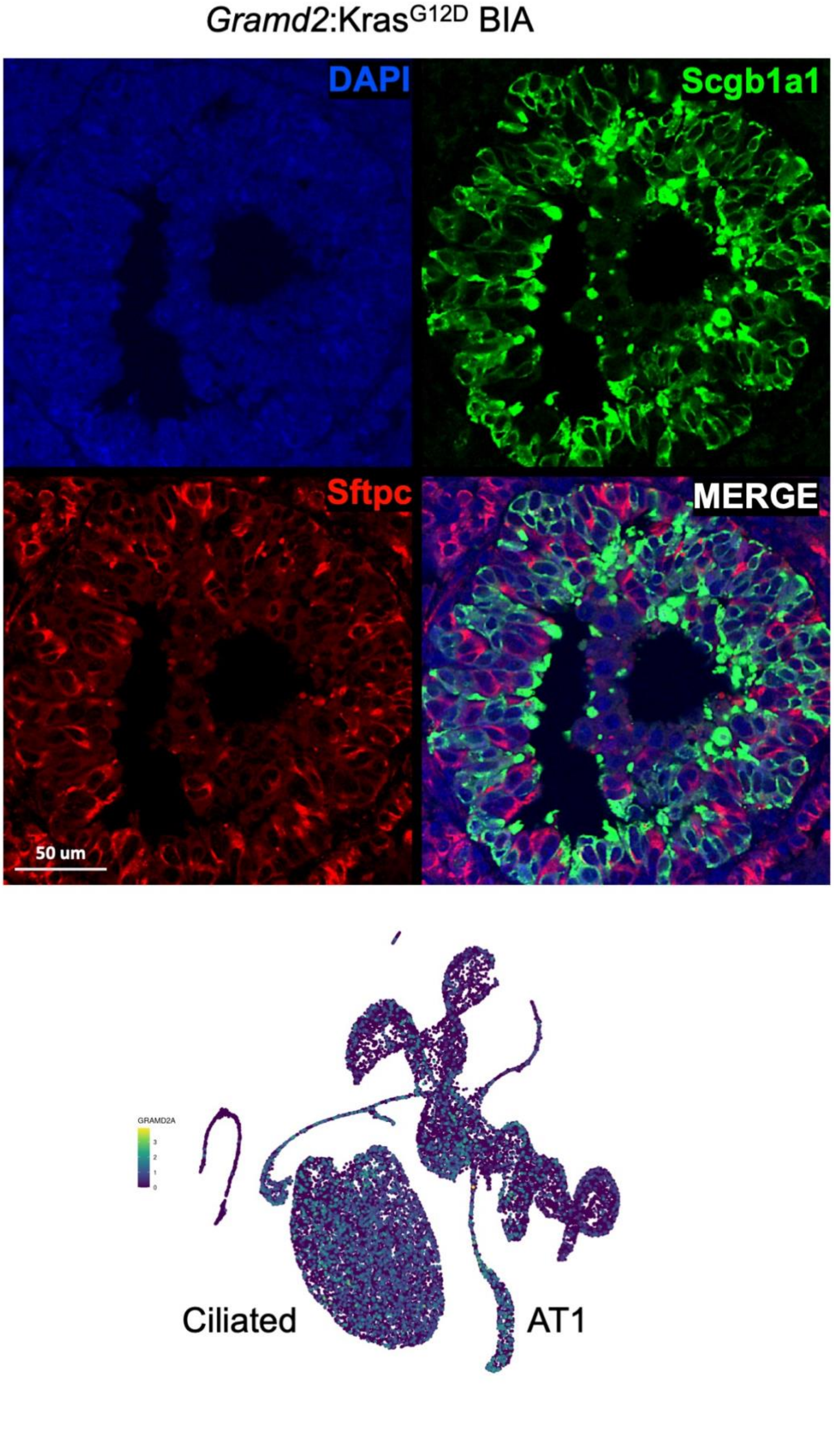
Gramd2:Kras$^{G12D}$ BIA
DAPI
Scgb1a1
Sftpc
MERGE
50 um
GRAMD2A
Ciliated
AT1

**Supplemental Figure 2: Bronchial infiltrative adenocarcinoma (BIA-LUAD) observed in *Gramd2:*Kras$^{G12D}$.** Top) IF staining of BIA lesion in *Gramd2:*Kras$^{G12D}$. Red = Sftpc (AT2 cell marker), Green = Scgb1a1 (club cell marker), blue = DAPI (nuclear stain). White bar = 50 um. Bottom) IPFCellAtlas expression of *GRAMD2A* (ipfcellatlas.com) (17).

**Figure S3**

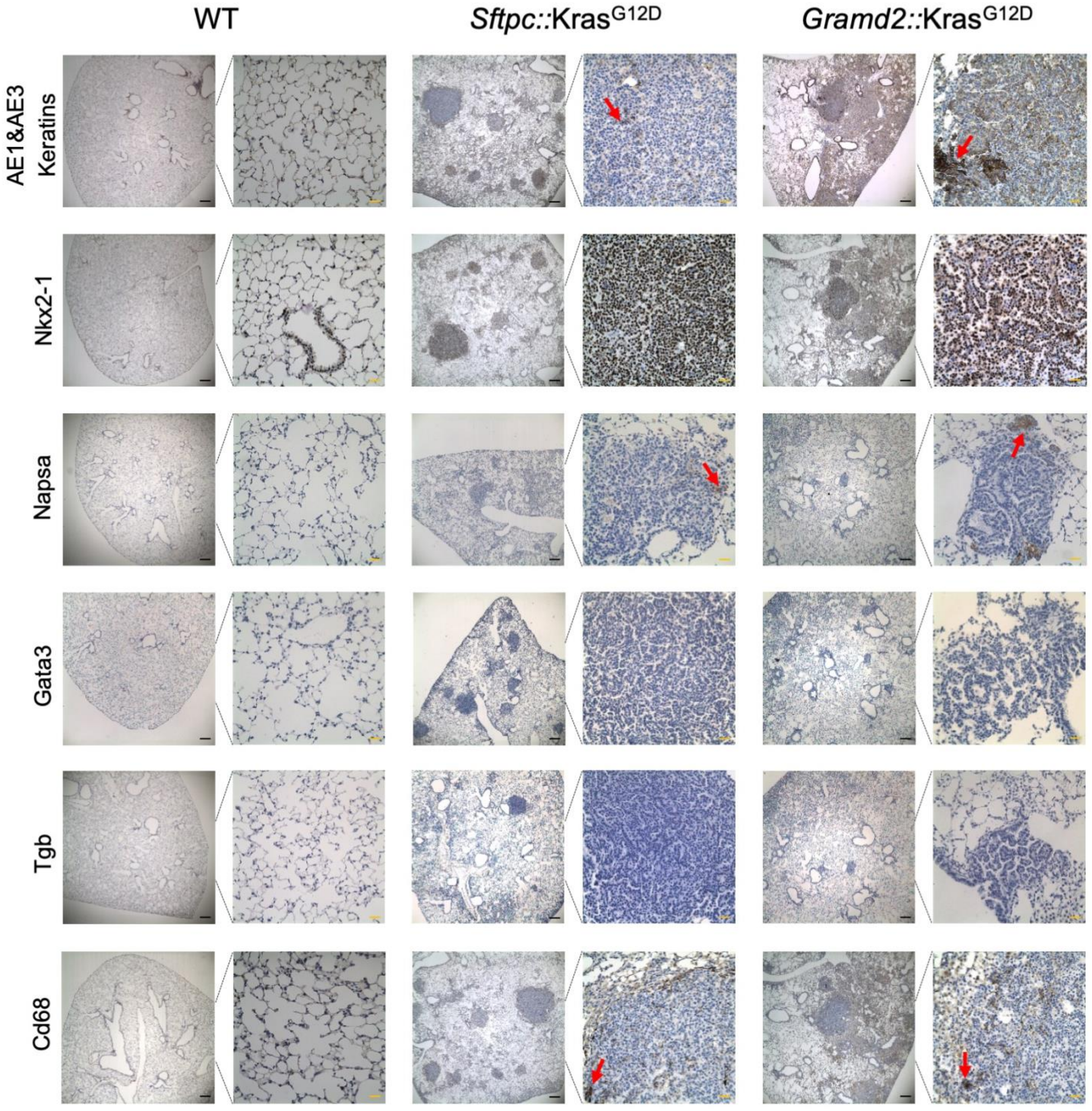

WT
*Sftpc::*Kras$^{G12D}$
*Gramd2::*Kras$^{G12D}$
AE1&AE3 Keratins
Nkx2-1
Napsa
Gata3
Tgb
Cd68

**Supplemental Figure 3: Immunohistochemistry staining profiles of *Gramd2*:Kras$^{G12D}$ mice are consistent with known staining patterns of papillary adenocarcinoma in human LUAD.** Staining for pan-cytokeratin AE1 & AE3 (Krt1-8, Krt10, Krt14-16 & Krt19)**,** Nkx2-1, Napsin A (Napsa), GATA3, Thyroglobulin (Tgb) and Cd68 on microphotographs of sections (4 μm) from WT, Kras$^{LSL-G12D}$, *Sftpc*-CreERT2, *Gramd2*-CreERT2, *Sftpc*:Kras$^{G12D}$ and *Gramd2*:Kras$^{G12D}$ mouse lungs 14 weeks after tamoxifen treatment (scale bar in black = 100 μm; scale bar in orange = 25 μm.). N= 3 per genotype.

### Figure S4

#### *Sftpc*:Kras$^{G12D}$ LUNG # 1

1,320
Number of Spots Under Tissue
144,242
Mean Reads per Spot
6,800
Median Genes per Spot

| | |
|---|---|
| Reads Mapped to Probe Set | 95.6% |
| Reads Mapped Confidently to Probe Set | 93.1% |
| Reads Mapped Confidently to the Filtered Probe Set | 86.0% |
| Number of Reads | 190,399,366 |
| Valid Barcodes | 96.9% |
| Valid UMIs | 99.7% |
| Sequencing Saturation | 79.8% |
| Q30 Bases in Barcode | 96.1% |
| Q30 Bases in Probe Read | 88.7% |
| Q30 Bases in UMI | 96.5% |

#### *Gramd2*:Kras$^{G12D}$ LUNG # 1

2,068
Number of Spots Under Tissue
178,867
Mean Reads per Spot
6,755
Median Genes per Spot

| | |
|---|---|
| Reads Mapped to Probe Set | 97.4% |
| Reads Mapped Confidently to Probe Set | 94.1% |
| Reads Mapped Confidently to the Filtered Probe Set | 86.3% |
| Number of Reads | 369,897,887 |
| Valid Barcodes | 98.2% |
| Valid UMIs | 99.7% |
| Sequencing Saturation | 83.2% |
| Q30 Bases in Barcode | 96.1% |
| Q30 Bases in Probe Read | 89.1% |
| Q30 Bases in UMI | 96.5% |

#### *Sftpc*:Kras$^{G12D}$ LUNG # 2

1,588
Number of Spots Under Tissue
159,806
Mean Reads per Spot
6,930
Median Genes per Spot

| | |
|---|---|
| Reads Mapped to Probe Set | 97.5% |
| Reads Mapped Confidently to Probe Set | 95.5% |
| Reads Mapped Confidently to Filtered Probe Set | 88.3% |
| Number of Reads | 253,771,413 |
| Valid Barcodes | 98.1% |
| Valid UMIs | 99.7% |
| Sequencing Saturation | 80.9% |
| Q30 Bases in Barcode | 96.2% |
| Q30 Bases in Probe Read | 89.0% |
| Q30 Bases in UMI | 96.6% |

#### *Gramd2*:Kras$^{G12D}$ LUNG # 2

2,083
Number of Spots Under Tissue
137,010
Mean Reads per Spot
4,269
Median Genes per Spot

| | |
|---|---|
| Reads Mapped to Probe Set | 94.6% |
| Reads Mapped Confidently to Probe Set | 74.8% |
| Reads Mapped Confidently to the Filtered Probe Set | 62.9% |
| Number of Reads | 285,390,830 |
| Valid Barcodes | 97.3% |
| Valid UMIs | 99.7% |
| Sequencing Saturation | 86.8% |
| Q30 Bases in Barcode | 96.1% |
| Q30 Bases in Probe Read | 88.5% |
| Q30 Bases in UMI | 96.5% |

**Supplemental Figure 4: Sequencing statistics for Visium 10X spatial transcriptomic analysis of 2 replicates of *Sftpc*:Kras$^{G12D}$ and *Gramd2*:Kras$^{G12D}$**. Q30 = quality score of greater than 30 per base, UMI = unique molecular identifiers.

### Figure S5

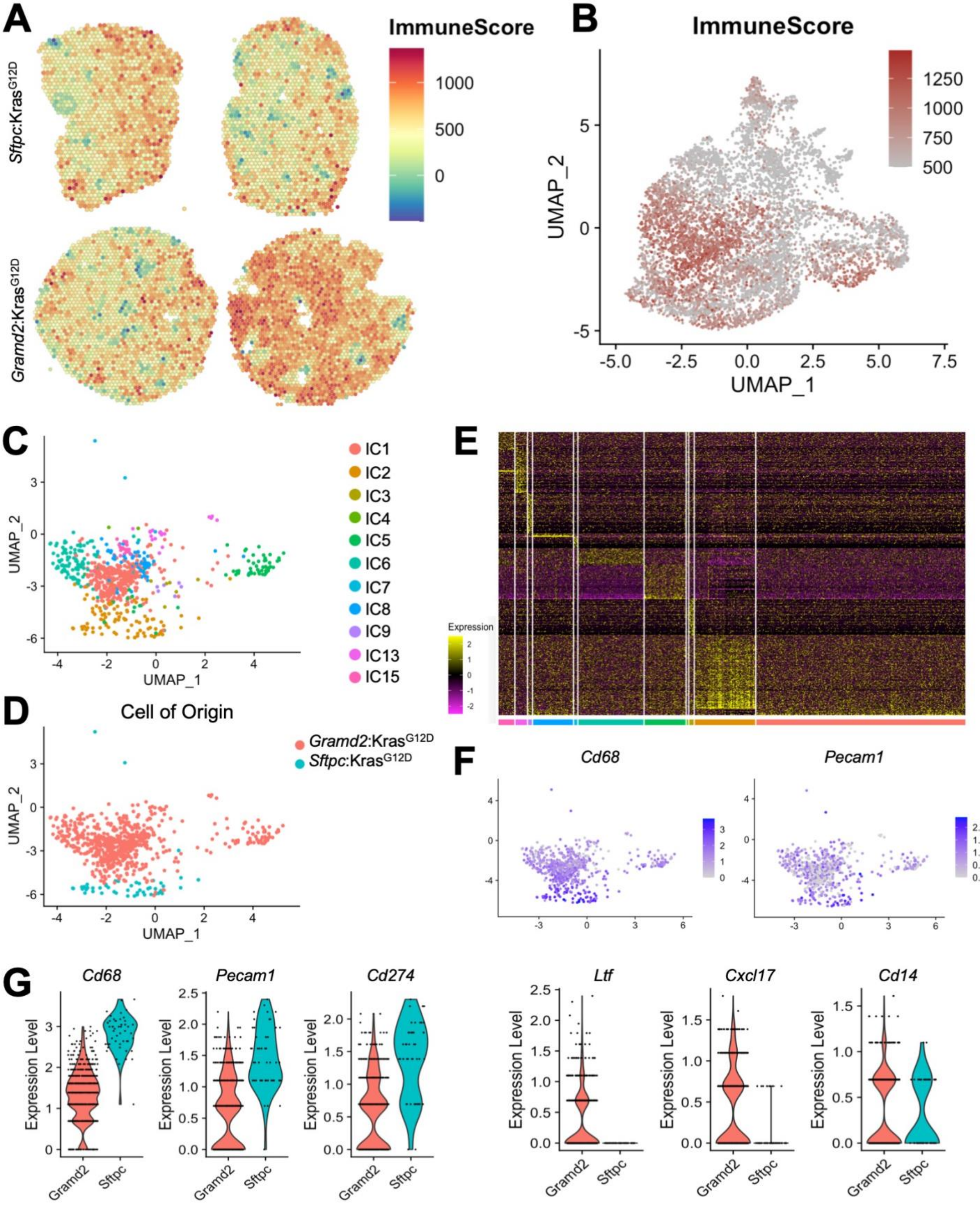

A
ImmuneScore
Sftpc:Kras$^{G12D}$
Gramd2:Kras$^{G12D}$
B
ImmuneScore
UMAP_1
UMAP_2
C
IC1
IC2
IC3
IC4
IC5
IC6
IC7
IC8
IC9
IC13
IC15
D
Cell of Origin
Gramd2:Kras$^{G12D}$
Sftpc:Kras$^{G12D}$
E
Expression
F
Cd68
Pecam1
G
Cd68
Pecam1
Cd274
Ltf
Cxcl17
Cd14
Expression Level
Gramd2
Sftpc

**Supplemental Figure 5: ESTIMATE analysis reveals differential immune cell infiltration between *Gramd2*:Kras$^{G12D}$ and *Sftpc*:Kras$^{G12D}$ driven LUAD.** A) ESTIMATE was applied to quantitate immune composition within samples; red = high immune cell signature, blue = low immune cell signature. B) UMAP projection of integrated clusters between all samples; brown = high immune cell signature, gray = low immune cell signature. C) UMAP projection of Visium 10X spots for all samples with an immune score >1000. Colors as indicated for Integrated Clusters (ICs). D) UMAP projection for array spots with immune scores >1000. Colors indicate sample origin; *Gramd2*:Kras$^{G12D}$ = pink, *Sftpc*:Kras$^{G12D}$ = teal. E) Heatmap of top 200 differentially expressed genes in ICs enriched for immune score. Yellow = high gene expression, purple = low gene expression. IC color as in (C). F) UMAP projection for array spots with immune scores >1000. Colors indicate expression *Cd68* (left) or *Pecam1* (right), purple = elevated expression, gray = little to no expression. G) Violin plots of select immune genes significantly differentially expressed between cells of origin. Expression is stratified by cell of origin; *Gramd2*:Kras$^{G12D}$ = pink, *Sftpc*:Kras$^{G12D}$ = teal.

### Figure S6

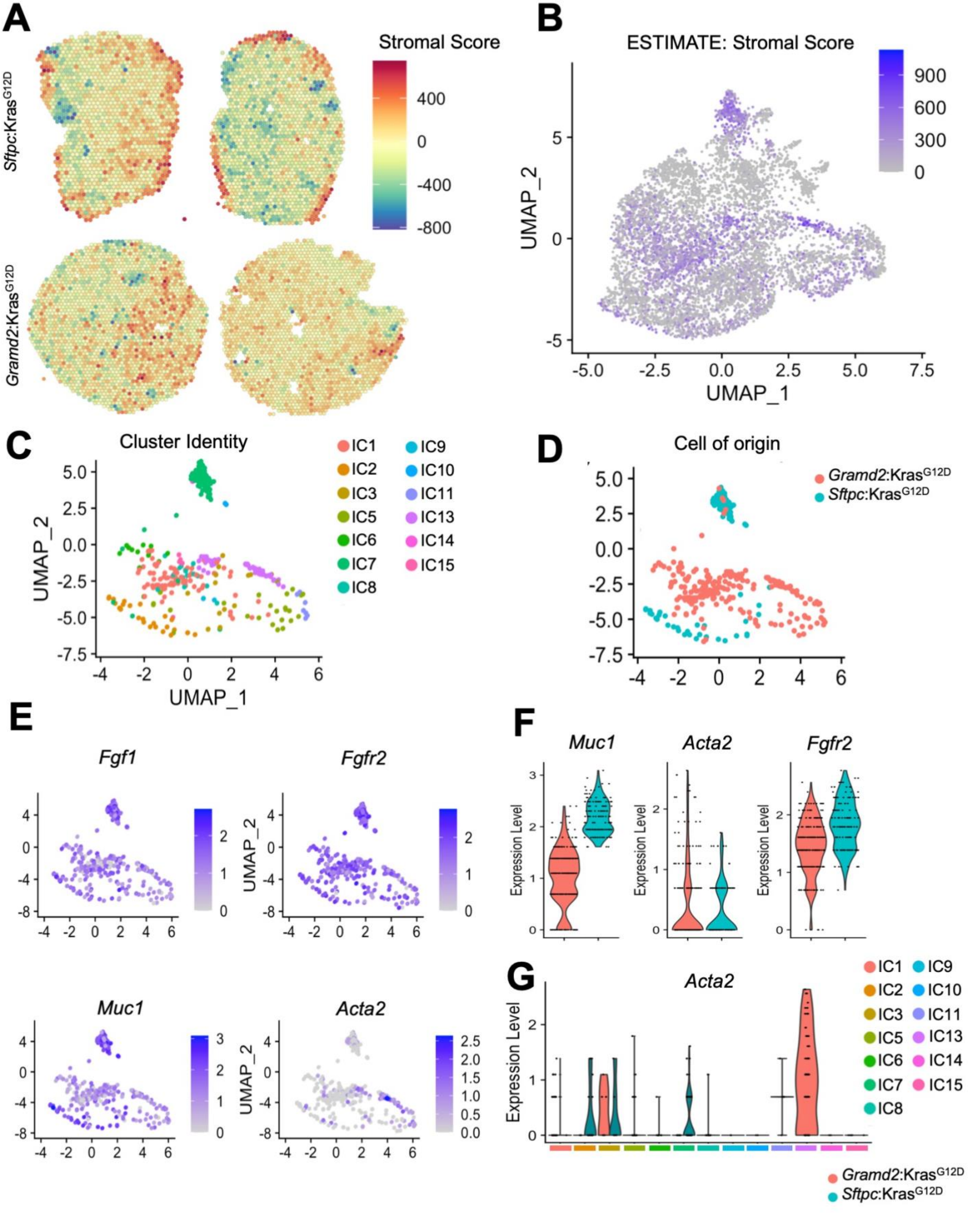
A
Stromal Score
Sftpc:Kras^G12D
Gramd2:Kras^G12D
B
ESTIMATE: Stromal Score
UMAP_1
UMAP_2
C
Cluster Identity
IC1 IC2 IC3 IC5 IC6 IC7 IC8 IC9 IC10 IC11 IC13 IC14 IC15
D
Cell of origin
Gramd2:Kras^G12D
Sftpc:Kras^G12D
E
Fgf1
Fgfr2
Muc1
Acta2
F
Muc1
Acta2
Fgfr2
Expression Level
G
Acta2
Expression Level
Gramd2:Kras^G12D
Sftpc:Kras^G12D

**Supplemental Figure 6: ESTIMATE analysis reveals differential stromal gene signatures between *Gramd2*:Kras$^{G12D}$ and *Sftpc*:Kras$^{G12D}$ driven LUAD.** A) ESTIMATE was applied to quantitate stromal composition within samples; red = high stromal gene signatures, blue = low stromal gene signatures. B) UMAP projection of integrated clusters between all samples; purple = high stromal signatures, gray = low stromal signatures. C) UMAP projection of Visium 10X array spots for all samples with a stromal score of >600. Colors as indicated for Integrated Clusters (ICs). D) UMAP projection for array spots with stromal scores > 600. Colors as indicated for Integrated Clusters (ICs). *Gramd2*:Kras$^{G12D}$ = pink, *Sftpc*:Kras$^{G12D}$ = teal. E) UMAP projection for array spots with stromal scores≥ 600. Colors indicate expression of known stromal genes. Purple = elevated expression, gray = little to no expression. F) Violin plots of select stromal genes significantly differentially expressed between cells of origin. Expression is stratified by cell of origin; *Gramd2*:Kras$^{G12D}$ = pink, *Sftpc*:Kras$^{G12D}$ = teal. G) Violin plot of alpha-SMA (*Acta2*) gene expression. Expression is separated by cell of origin *Gramd2*:Kras$^{G12D}$ = pink, *Sftpc*:Kras$^{G12D}$ = teal, as well as by IC grouping. Colors as indicated for Integrated Clusters (ICs).

### Figure S7

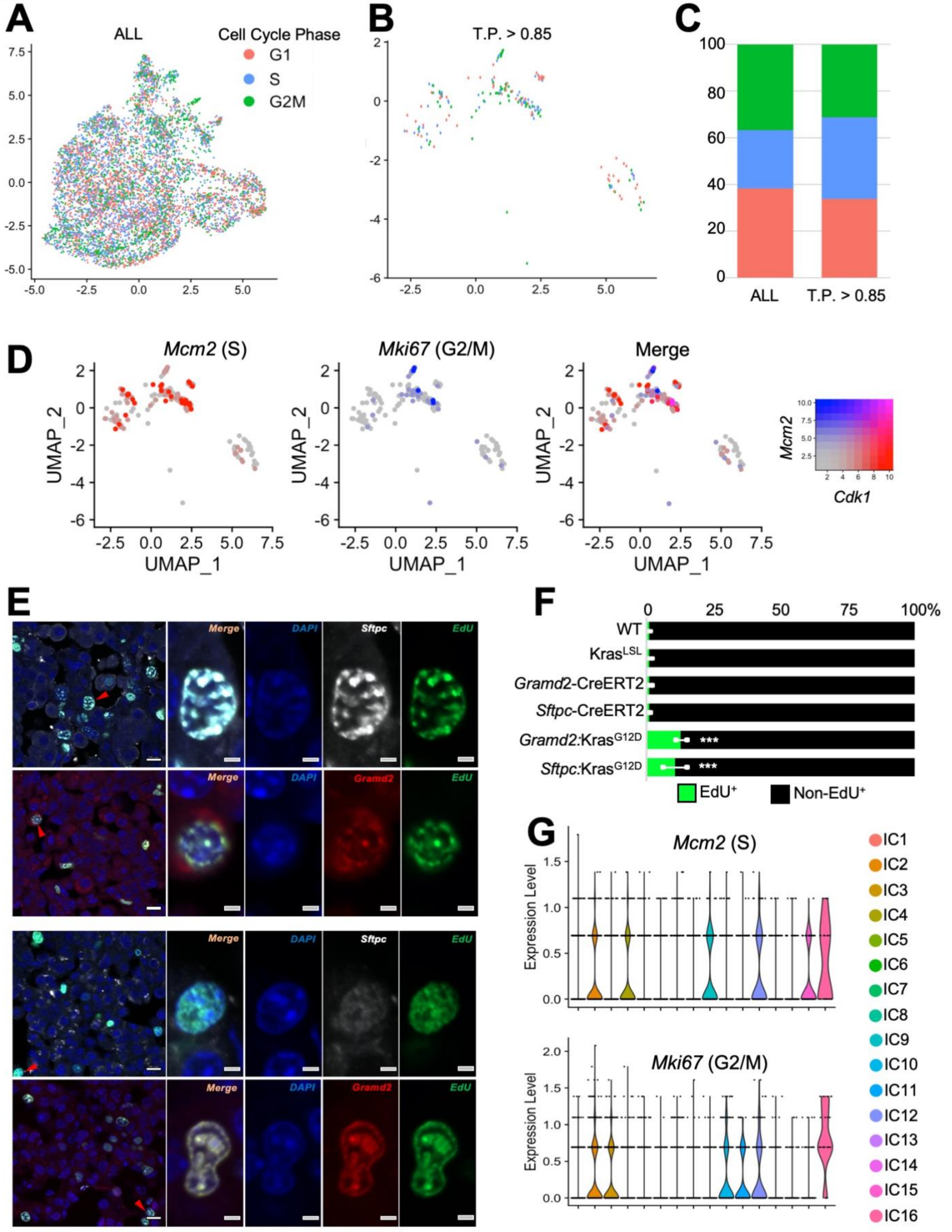
A
ALL
Cell Cycle Phase
G1
S
G2M
B
T.P. > 0.85
C
ALL
T.P. > 0.85
D
Mcm2 (S)
Mki67 (G2/M)
Merge
UMAP_1
UMAP_2
Mcm2
Cdk1
E
Sftpc:KrasG12D
Gramd2:KrasG12D
Merge
DAPI
Sftpc
Gramd2
EdU
F
WT
KrasLSL
Gramd2-CreERT2
Sftpc-CreERT2
Gramd2:KrasG12D
Sftpc:KrasG12D
***
EdU+
Non-EdU+
G
Mcm2 (S)
Mki67 (G2/M)
Expression Level
IC1
IC2
IC3
IC4
IC5
IC6
IC7
IC8
IC9
IC10
IC11
IC12
IC13
IC14
IC15
IC16

**Supplemental Figure 7: LUAD derived from both *Sftpc*:Kras$^{G12D}$ and *Gramd2*:Kras$^{G12D}$ exhibit elevated proliferation rates.** A) UMAP projection of Integrated Clusters between all (ALL) samples; red = Visium 10X array spots in G0 or G1 phase (the absence of S or G2/M signatures), blue = array spots with cells in S phase, green = array spots with cells in G2 or M phases. B) UMAP projection of Visium 10X spots for all samples with tumor purity ≥ 0.85 (85%). Colors = G0/G1 (red), S (blue), G2/M as in (A). C) Proportion of Visium 10X array spots in each phase of the cell cycle between all spots (ALL) and those with Tumor Purity ≥ 0.85. D) Occurrence of *Mcm2* (S phase marker) and *Mki67* (G2/M phase marker) in Visium array spots with Tumor Purity >0.85. Red = *Mcm2* expressing blue = *Mki67* expressing, purple = co-expression of both *Mcm2* and *Mki67* within the same array spot. E) Immunofluorescence performed on *Sftpc*:Kras$^{G12D}$ (top) and *Gramd2*:Kras$^{G12D}$ (bottom) samples in histologically defined LUAD regions. DAPI = blue, Sftpc = white, Gramd2 = red, EdU incorporation = green. N = 3 per genotype. F) Quantification of EdU+) nuclei expressed as a percent of total DAPI$^{+}$ nuclei. N = 3 for each genotype. ANOVA Significance *** <0.001. G) Violin plot of *Mcm2* (S phase marker) and *Mki67* (G2/M phase marker) expression within each Integrated Cluster (IC) present across the entire sample set.

### Figure S8

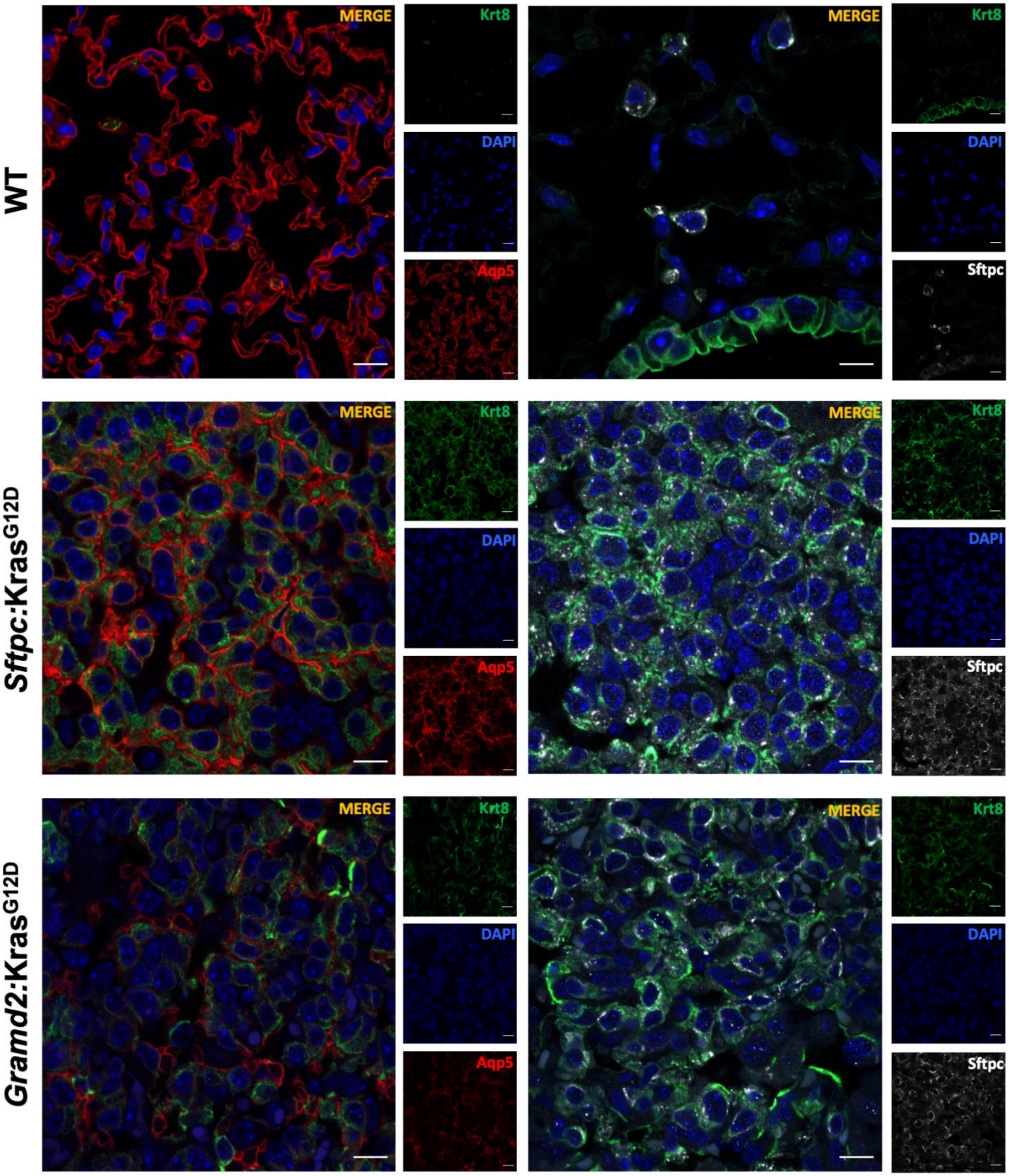
WT
Sftpc:Kras$^{G12D}$
Gramd2:Kras$^{G12D}$
MERGE
Krt8
DAPI
Aqp5
Sftpc

**Supplemental Figure 8. Higher magnification of Krt8$^+$ staining in AT2 and AT1 cell-derived LUAD**. WT = C57BL/6 control mice, *Sftpc:*Kras$^{G12D}$ and *Gramd2*:Kras$^{G12D}$ mouse lung (scale bar is 10 µm). Red = Aqp5 (AT1 cell marker), white = Sftpc (AT2 cell marker), green = Krt8 (AEC-intermediate / transitional cell state marker), blue = DAPI (nuclear stain). Scale (white bar) = 10 µM (63x).

**Figure S9**

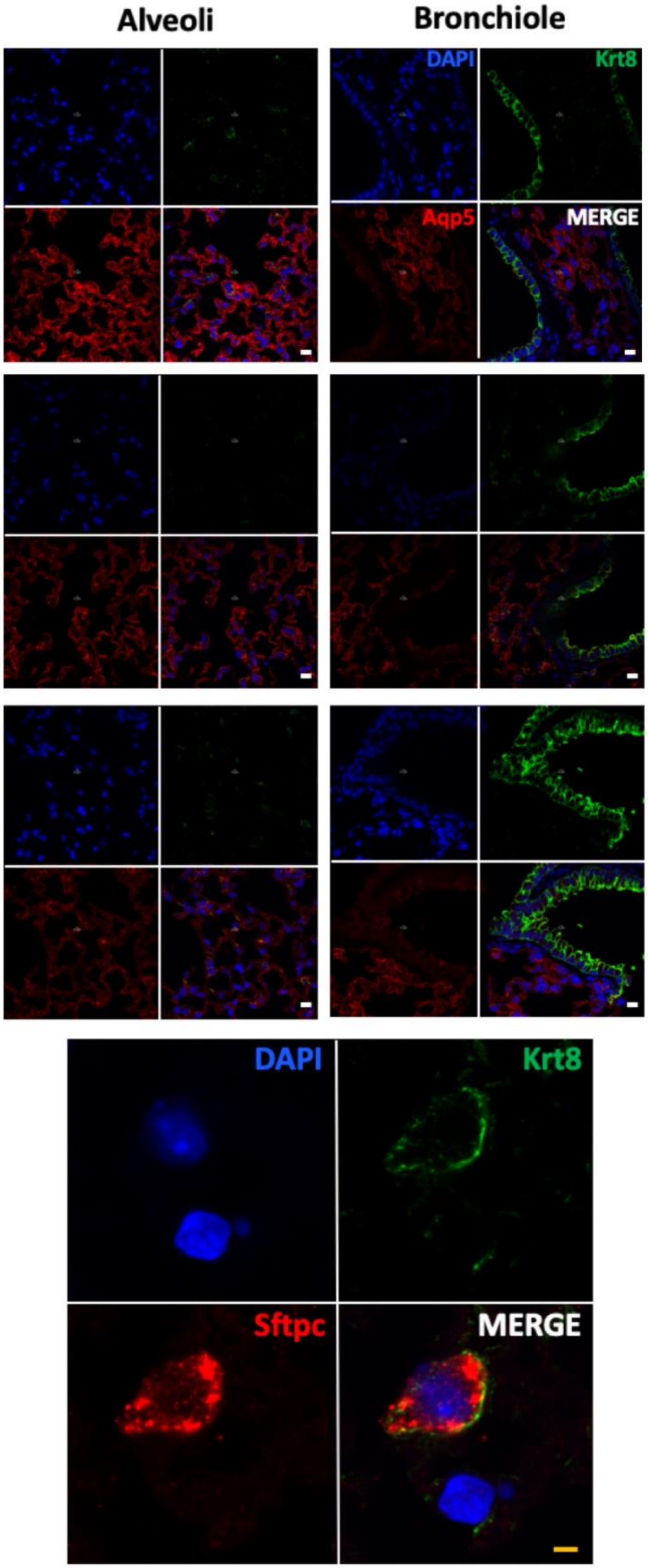

Alveoli
Bronchiole
DAPI
Krt8
Aqp5
MERGE
DAPI
Krt8
Sftpc
MERGE

**Supplemental Figure 9. Control Krt8 staining patterns in wild type (WT) mouse lungs**. Upper panels: WT = C57BL/6 control mice, *Sftpc*:Kras$^{G12D}$ and *Gramd2*:Kras$^{G12D}$ mouse lung (scale bar is 10 µm). Red = Aqp5 (AT1 cell marker), green = Krt8 (AEC-intermediate / transitional cell state marker), blue = DAPI (nuclear stain). Lower panel: High magnification of red = Sftpc (AT2 cell marker), green = Krt8 (AEC-intermediate / transitional cell state marker), blue = DAPI (nuclear stain) in WT C57BL/6. Scale (white bar) = 20 µM (20x); scale (orange bar) = 10 µM (63x).
